# Two serial filters determine P2X7R cation selectivity, Ser342 in the central pore and lateral acidic residues at the cytoplasmic interface

**DOI:** 10.1101/2024.02.19.579953

**Authors:** Fritz Markwardt, Eike Schön, Michaela Raycheva, Aparna Malisetty, Sanaria Hawro Yakoob, Malte Berthold, Günther Schmalzing

## Abstract

The human P2X7R (hP2X7R) is a homotrimeric cell surface receptor gated by extracellular ATP^4-^ with two transmembrane helices per subunit, TM1 and TM2. A ring of three S342 residues, one from each pore-forming TM2 helix, located halfway across the membrane bilayer, functions to close and open the gate in the apo and ATP^4-^ bound open states, respectively. The hP2X7R is selective for small inorganic cations, but can also conduct larger organic cations such as Tris^+^. Here, we show by voltage-clamp electrophysiology in *Xenopus laevis* oocytes that mutation of S342 residues to positively charged lysines decreases the selectivity for Na^+^ over Tris^+^, but maintains cation selectivity. Deep in the membrane, laterally below the S342 ring are nine acidic residues arranged as an isosceles triangle consisting of residues E14, D352, and D356 on each side, which do not move significantly during gating. When the E14K mutation is combined with lysine substitutions of D352 and/or D356, cation selectivity is lost and permeation of the small anion Cl^-^ is allowed. Lysine substitutions of S342 together with D352 or E14 plus D356 in the acidic triangle convert the hP2X7R mutant to a fully Cl^-^-selective ATP^4-^-gated receptor. We conclude that the ion selectivity of wild-type hP2X7R is determined by two sequential filters in one single pathway: (1) a primary size filter, S342, in the membrane center and (2) three cation filters lateral to the channel axis, one per subunit interface, consisting of a total of nine acidic residues at the cytoplasmic interface.

**Significance:** Pore size and electrostatic interactions are key to the permeation selectivity of ion channels. Previous cysteine scanning mutagenesis identified a tri-serine-342 ring located halfway across the membrane as the gate and selectivity filter of the P2X7 receptor channel, accessible from the inside to cationic but not anionic reagents. Consistent with a downstream cation filter, we could now switch P2X7R from cation to anion selectivity by lysine substitution of acidic residues at the cytoplasmic interface. Our data show that two sequential selectivity filters control the cation selectivity of the P2X7R channel, a dynamic tri-serine-342 size filter and three conformationally static cation filters of three acidic residues each. We propose that the ion selectivity of P2X receptors involves the mechanism described here.

## Introduction

The P2X7 receptor (P2X7R) is an ATP^4-^-gated cation-selective channel that is of great interest as a drug target (1) because of its widespread expression in virtually all immune cells, including microglia (2) and also tumor cells (3), its role in inflammation, and also because of its enigmatic ability to generate large cytotoxic pores (4). Extracellular binding of the ligand ATP^4-^ opens the channel pore within milliseconds to small cations such as Na^+^, K^+^, and Ca^2+^, but also allows permeation of larger organic cations, including Tris^+^ (121 Da), N-methyl-d-glucamine^+^ (NMDG, 195 Da) and cationic DNA-binding dyes such as ethidium^+^ (314 Da) and Yo-Pro-1^2+^ (376 Da).

For many years, the permeation of large cations was attributed to a second conducting state in which the P2X7R channel pore gradually increases upon continued stimulation with ATP (5), eventually leading to membrane blebbing and cytolysis (6, 7). In contrast to whole cell recordings, single-channel recordings of ATP^4-^-induced currents in hP2X7R-expressing human B lymphocytes (8) and *X. laevis* oocytes provided no evidence for a time-dependent increase in pore diameter (9). Instead, single-channel recordings showed a low instantaneous permeability to Tris^+^ (mean diameter 5.8 Å, for ionic diameter references see Table 1), NMDG^+^ (mean diameter 7.3 Å) and other large cations, which remained constant until ATP^4-^ was washed out (10). This view is supported and extended by data showing instantaneous robust NMDG^+^ currents in HEK293 cells with the P2X2 receptor (11) and all homotrimeric P2X receptors except P2X1 (12). Also the purified liposomal reconstituted panda P2X7R (pdP2X7R), which shares 85 % sequence identity with hP2X7R, has been shown to be intrinsically permeable to the cationic dye YO-PRO-1 (376 Da, mean diameter 12 Å) (13).

**Table 1.** Ion diameter compilation with references.

| Cation | Ion size (Å) | Mean diameter<sup>1</sup> (Å) | Reference |
| --- | --- | --- | --- |
| K<sup>+</sup> | 2.66 (dehydrated) |  | (60) |
| MTSEA<sup>+</sup> | 4.6 x 4.6 x 10 |  | (17) |
| MTSET<sup>+</sup> | 5.8 x 5.8 x 10 |  | (17) |
| Na<sup>+</sup> | 1.90 (dehydrated) |  | (60) |
| Na<sup>+</sup> | ~2.1 (dehydrated) |  | (61) |
| NMDG<sup>+</sup> | 5.0 x 6.4 x 12.0 | 7.3 | (29) |
| NMDG<sup>+</sup> | 6.5 x 8.7 x 10.8 | 8.4 | (62) |
| Tris<sup>+</sup> | 6.0 x 6.9 x 7.7 | 6.8 | (63) |
| Tris<sup>+</sup> | 5.5 x 5.6 x 6.4 | 5.8 | (29) |
| Tris<sup>+</sup> | 6.4 x 6.5 x 7.5 | 6.8 | (62) |
| Yo-Pro-1<sup>2+</sup> | 8.2 x 12.8 x 16.8 | 12.0 | (62) |
| **Anion** |  |  |  |
| Cl<sup>-</sup> | 3.62 |  | (60) |
| Glu<sup>-</sup> | 5.5 x 6.5 |  | (64) |
| MTSES<sup>-</sup> | 5.8 x 5.8 x 10 |  | (17) |
| **Zwitterion** |  |  |  |
| Hepes | ~10 Å |  | (65) |

<sup>1</sup>Mean diameter was calculated as the geometric mean of the dimensions of a box containing the ion, as previously proposed (29).

Plotting the single-channel conductance against the apparent molecular diameter of the permeating organic cations yielded an ATP^4-^-opened effective pore diameter of ∼8.5 Å of the hP2X7R, which was stable over time (10). Considering the different measurement methods (single channel recordings vs. cryo-EM) and species (human vs. rat), the open pore diameter of ∼8.5 Å in *X. laevis* oocytes is in reasonable agreement with the structural diameter of 5.0 Å at S342 of rP2X7R reconstituted in detergent micelles (14). The apo-closed cryo-EM structure of rP2X7R shows that the pore is completely collapsed between residues S339 and S342, which define the extracellular and intracellular boundaries of the TM2 gate, respectively (14). This is consistent with cysteine scanning accessibility mutagenesis, which showed that the closed channel is accessible from the outside up to residue S339, but not up to S342 (15).

A tri-serine ring with a pore diameter of 5.0 Å and larger, which might allow even fully hydrated Na^+^ or K^+^ with diameters of ∼5 Å (16) or ∼6 Å (17) to pass, is difficult to reconcile with the selective discrimination of small cations. We therefore asked here whether the hP2X7R possesses acidic residues that discriminate between anions and cations by repulsion and attraction. An exception to the strict cation permeability of other members of the P2XR family is the P2X5R, which exhibits significant anion permeability in addition to cation permeability (18, 19). Recently, a basic residue in P2X5 receptors that replaces glutamate in the cytoplasmic lateral fenestrations of purely cation-selective P2XRs was shown to be responsible for anion permeability (20). To investigate which residues in the transmembrane pathway of hP2X7R control the size and charge of permeant cations, we electrophysiologically characterized lysine mutants of S342 and downstream acidic residues at the plasma membrane-cytoplasmic interface and used homology models based on the apo-closed and open cryo-EM structure of hP2X7R (14) for interpretation. We found that cationic selectivity is determined by two serial filters in a single pathway, a primary size-screening filter midway in the membrane formed by a ring of three S342 residues, one per subunit, and a primary electrostatic filter further downstream and lateral to S342 formed by a total of 9 acidic residues, 3 per subunit, located in the membrane-cytoplasmic interface.

## Results

### In silico identification of candidate hP2X7R residues involved in cation selectivity

Like other functional P2XR isoforms (21, 22), the hP2X7R^wt^ assembles as a homotrimer in *X. laevis* oocytes (23). Fig. 1A shows a lateral view of a SWISS homology model of the apo-closed homotrimeric hP2X7R^wt^ based on the apo rP2X7R cryo-EM structure (14) with a large extracellular ectodomain containing intersubunit ATP^4-^-binding sites, a membrane-spanning domain consisting of two transmembrane α-helices per monomer, TM1 and TM2, and a large intracellular domain termed the cytoplasmic ballast (14). The TM1 helices (light gray) face outward. The three central TM2 helices (yellow) form a trihelical bundle that lines the transmembrane pore.

**Fig. 1.**
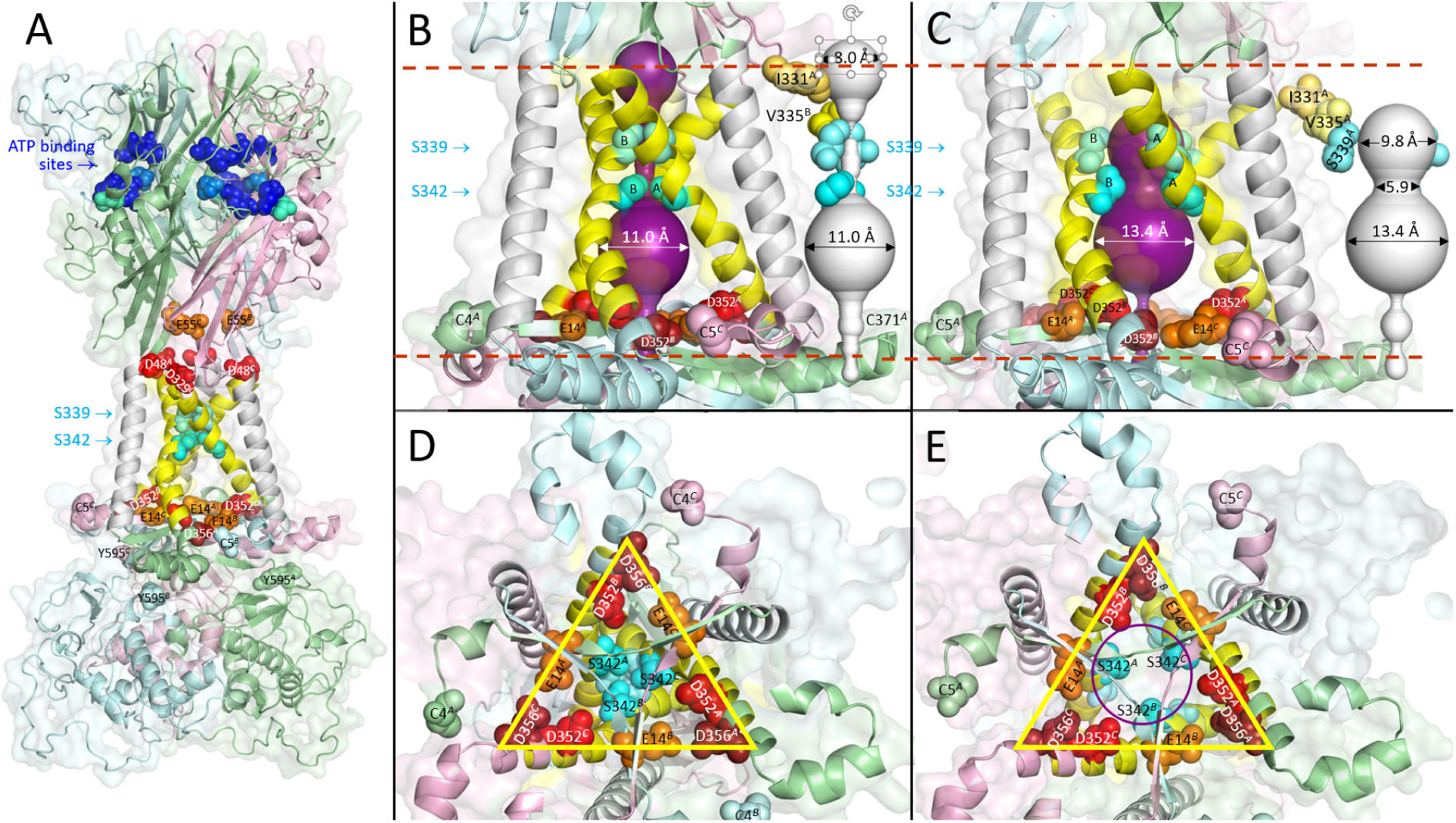
Visualization of candidate residues involved in controlling the cation selectivity of the hP2X7R. A, Overall structure of the hP2X7R^wt^ homotrimer homology-modeled based on the cryo-EM structure of the apo-closed conformation of the rP2X7R (14), viewed perpendicular to the membrane normal. The hP2X7R^wt^ extends over *∼*71 Å (N302^B^ to D329^C^) and *∼*50 Å (D356^B^ to S584^B^) in the extracellular and intracellular directions, with a maximum width of *∼*71 Å (G418^B^ to S290^A^). Each of the three identical subunits (*ABC*) is shown in a different color (A, pale-green, B pale-cyan, C light-pink), except for TM1 and TM2, which are shown in white and yellow, respectively, for easier identification. The three TM2 domains line the channel pore. Several residues are highlighted as spheres: C4 and C5, the first residues solved in the apo and open rP2X7R cryo-EM structures, respectively, and the last residues Y595 in their chain colors; S339 and S342 around the gate in aquamarine and cyan, respectively; in the ectodomain, the acidic residues in red and the residues that coordinate the ectodomain ATP^4-^ binding by ionic interactions (K64, K66, R294, K311) and hydrogen bonding (T189, N292) in blue and marine, respectively. B,C Enlarged lateral views of the transmembrane region of the closed and open states of hP2X7R^wt^, including the channel pore as predicted by MOLEonline (25). Channels are shown in duplicate, with surrounding protein (purple) and, for a free view of the channel shape, in white, with only the selected residues (one or two per triplet) accessible to cysteine-reactive reagents from the extracellular space: I331 (yellow-orange), Y335 (pale-yellow), S339 (aquamarine), and S342 (cyan). The dashed red lines indicate the outer and cytoplasmic boundaries of the membrane as predicted by the OPM database (24). The open channel model (C) lacks a sufficiently wide opening to the cytoplasm for ions to exit, as does the closed channel model (B). D,E Cytoplasmic views on the channel pore and surrounding acidic residues (E14 in orange, D352 in red, D356 in firebrick) at the level of the membrane-cytoplasmic interface after masking of the cytoplasmic structure. Obviously, residues S339 and S342 move laterally to open the pore (compare D to E). In contrast, the acidic residues arranged in an isosceles triangle (yellow) do not move significantly during channel opening (compare D to E).

In the ’pore’ and ’membrane ON’ modes, where the position of the lipid bilayer is included in the calculation according to the Orientations of Proteins in Membranes (OPM) database (24), the channel prediction tool MOLEonline (25) identifies only a short continuous intramembrane channel pore along the central axis of P2X7R^wt^ in both the apo-closed and ATP-bound open states (Fig. 1B and C), i.e. without any significant access from the outside or exit to the cytoplasm (for calculations without membrane constraint, see *SI Appendix*, Fig. S1). Enlarged views of the TM domains show that residues S339 and S342 (aquamarine and cyan spheres), one per TM2 helix, move laterally to effectively close (Fig. 1B, D) or open (Fig. 1C, E) the channel. The extracellular accessibility of I331 (yellow-orange), V335 (pale-yellow), and S339, but not S342, of the closed hP2X7R (Fig. 1B), as shown by cysteine scanning in single-channel and fluorescent dye binding measurements (15), is consistent with the homology model derived from the cryo-EM structures of rP2X7R (14).

Bounded at the top by S342 in both the closed and open states, there is a voluminous cavity in the second half of the membrane, followed further downstream by an extension into the cytoplasm formed by a ring of three centrally located basic K17 residues that is far too narrow to allow cations to exit into the cytoplasm (Fig. 1B, C, see also *SI Appendix*, Fig. S2). However, lateral to K17 are nine acidic residues, three per subunit (E14 and D352/D356 each from neighboring subunits), arranged in an isosceles triangle parallel to the membrane to form a massive negative charge cluster that could represent cation selectivity filters (Fig. 1D,E; see also *SI Appendix*, Fig. S2). Upon ATP^4-^ binding, the channel opens wide due to the lateral movement of S339 and S342, and the voluminous cavity further dilates somewhat, while the lateral nine acidic residues do not move significantly (compare Fig. 1 B,D and C,E; *SI Appendix*, Fig. S2).

### Like hP2X7R^wt^, the gate mutant hP2X7R^S342K^ is cation-selective

The principle of our ion selectivity measurements is shown in Fig. 2. I-V curves were generated during ATP-induced receptor activation in superfusion solutions containing Na^+^Cl^-^ (Fig. 2A-C), Tris^+^Cl^-^ (Fig. 2D-F), or Tris^+^Glu^-^ (Fig. 2G-I) as major ions. The intersection of the I-V curve with the zero current x-axis yields the reversal potential V_rev_, which is the potential at which the net current through the open channel, is zero. Replacing Na^+^ (ionic diameter 1.9-2.1 Å) with the three times larger organic cation Tris^+^ (diameter ∼6.4 Å) drastically shifted V_rev_ to the left from -6.8 mV (Fig. 2A) to -53.9 mV (Fig. 2D) in hP2X7R^wt^-expressing oocytes. This indicates that most of the current in Fig. 2A was carried by the small cation Na^+^ and that hP2X7R^wt^ has a much lower permeability for the larger cation Tris^+^. The calculated Tris^+^:Na^+^ permeability ratio P_Tris_:P_Na_ according to Hille (26) (see eq. 1) of 0.16 is in good agreement with our previous estimates from single channel recordings of 0.10 and 0.075 for native hP2X7R in human B lymphocytes (8) and recombinant hP2X7R in *X. laevis* oocytes (10), respectively.

**Fig. 2.**
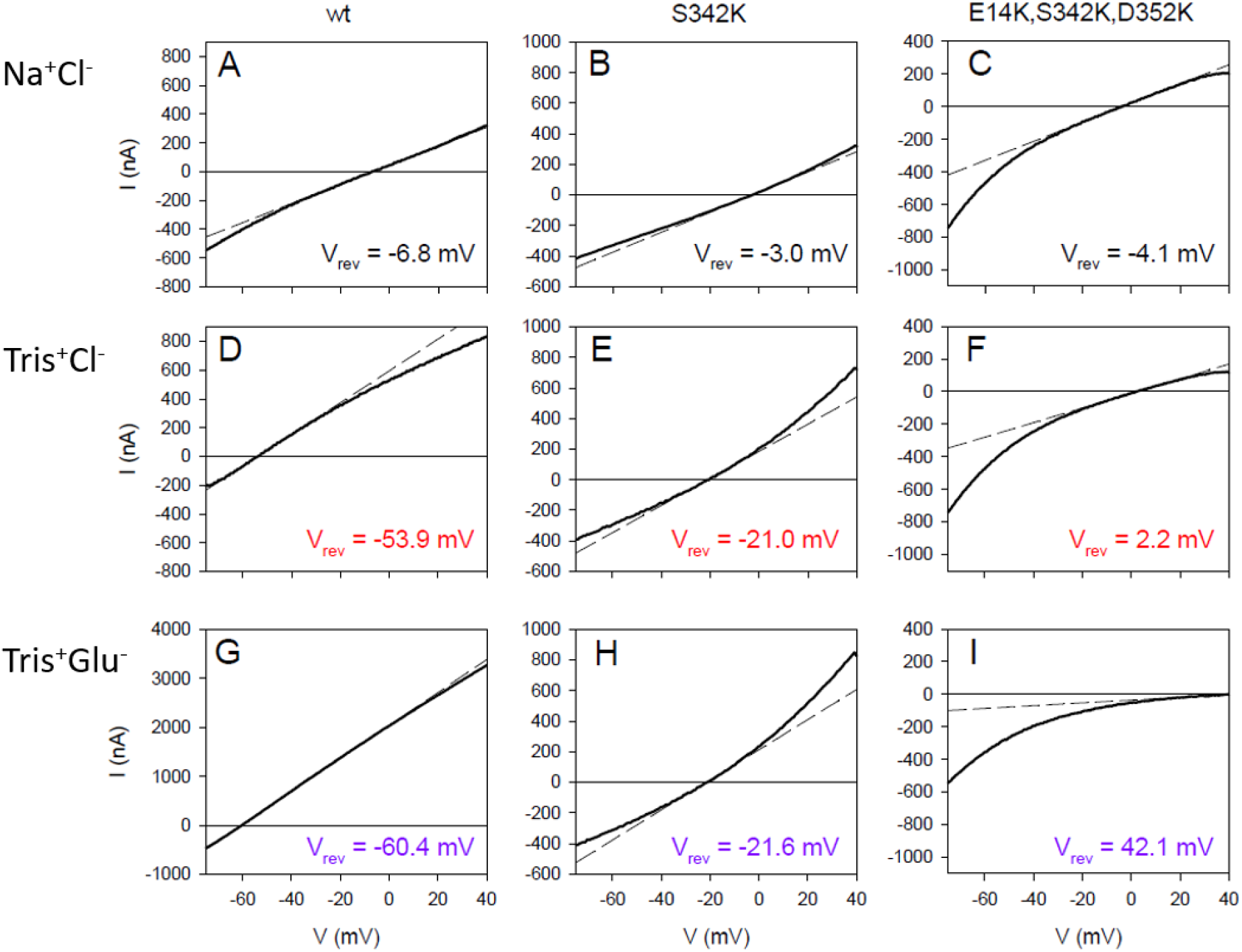
Ramp currents mediated by hP2X7R^wt^ and mutants expressed in *X. laevis* oocytes. The ATP-induced ramp currents shown were calculated as the difference between the ramp currents before and during 0.1 mM ATP^-4^ application. Only the ramp currents in the voltage range of -70 to +40 mV are shown because capacitive currents are generated by voltage jumps from holding potentials of -40 to -80 mV. The name of the hP2X7R construct expressed is shown at the top of each of the three rows of panels, and the only salt (except ∼5 mM Hepes, diameter ∼10 Å) of the extracellular solution is shown at the left margin. The bold curved lines show the measured currents; the thin dashed straight lines show the linear fit of the current-voltage relationship near the reversal potential. From this approximation, the V_rev_ values (shown in each of the figures here and in Fig. 3) and the conductance (slope of the lines whose statistics are shown as G_at Vrev_ in Fig. 3) were calculated.

Additional substitution of extracellular Cl^-^ (ionic diameter 3.6 Å) by the almost 2-fold larger organic anion Glu^-^ (dimensions 5.5 x 6.5 Å) to test for anion selectivity induced only a slight shift of V_rev_ to more negative potentials from -53.9 mV in Tris^+^Cl^-^ (Fig. 2D) to -60.4 mV in Tris^+^Glu^-^ (Fig. 2G; not significant, for statistics see Fig. 3). For an anion-permeable ion channel, one would have expected an opposite shift to more positive reversal potentials (see Fig. 2F and I). The current result is consistent with previous findings that native and recombinant hP2X7R^wt^ are Cl^-^-impermeable in physiological media (8, 27).

**Fig. 3.**
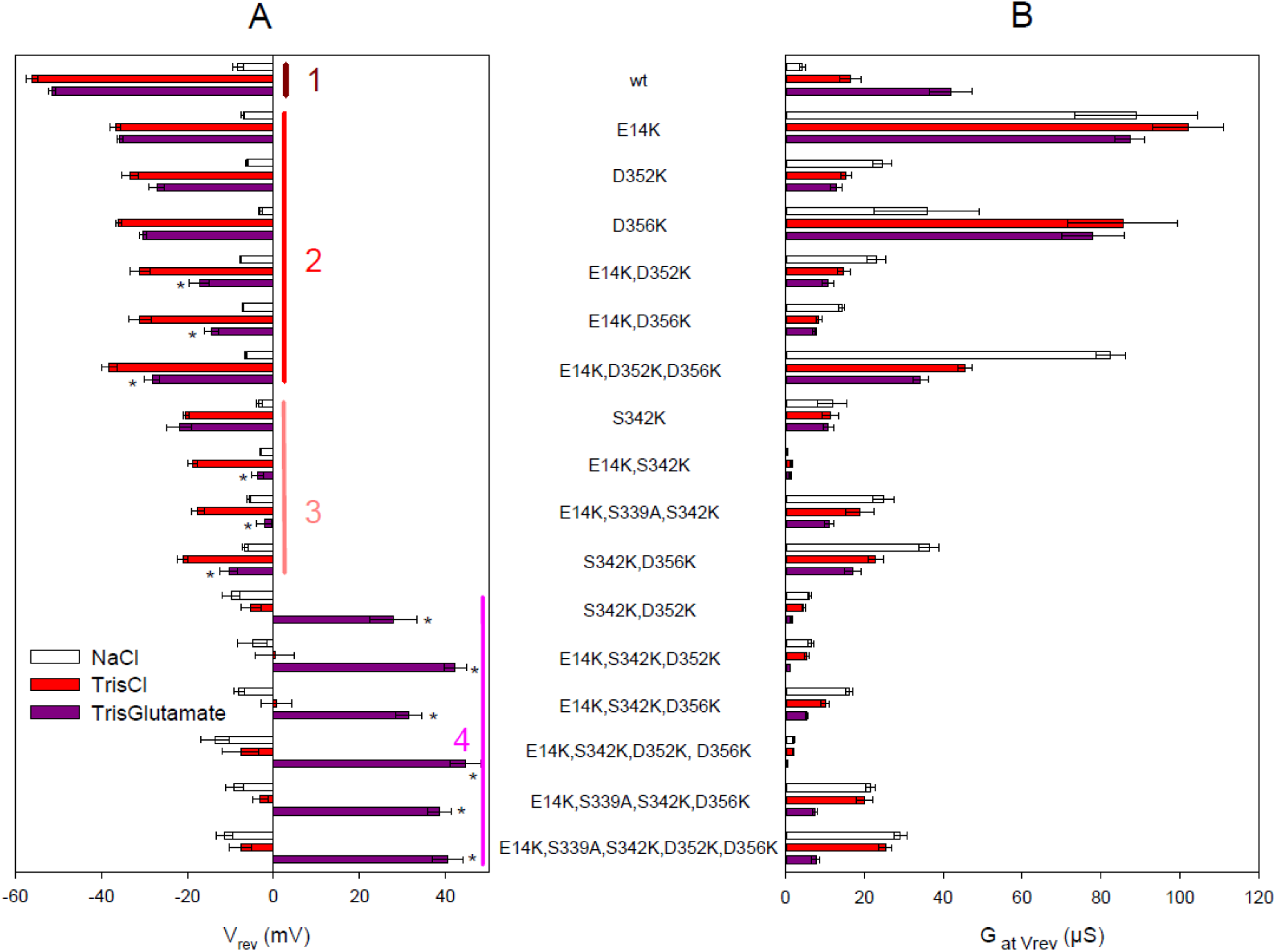
Permeation characteristics of hP2X7R constructs expressed in *X. laevis* oocytes. Bars represent the reversal potential (A) and the corresponding slope conductance at V_rev_ (B) of the indicated hP2X7R constructs during application of 0.1 mM ATP^-4^ in one of three extracellular solutions based on Na^+^Cl^-^ (open columns), Tris^+^Cl^-^ (red columns), and Tris^+^Glu^-^ (purple columns). Data are mean ± SEM of 6-7 oocytes. (A) The constructs are arranged in 4 correspondingly numbered groups that differ significantly in their reversal potential in Tris^+^Cl^-^. Group 1: hP2X7R^wt^. Group 2: Substitution of single or multiple acidic residues by lysine as indicated; note that the V_rev_ shift due to substitution of Na^+^ by Tris^+^ (i.e. the difference between the white and red columns) of all mutants in this group is significantly smaller than that of hP2X7R^wt^, indicating that they all have reduced selectivity for small cations. Group 3: Note that the S^342^K mutation further reduces the V_rev_ shift in Tris^+^Cl^-^ vs. NaCl compared to the mutants in group 2, indicating a further reduced selectivity for the small cation Na^+^; the additional mutation of E14 or D356 to lysine additionally produces a significant V_rev_ shift in Tris^+^Glu^-^ vs. Tris^+^Cl^-^, most likely reflecting an occurring anion permeability with reduced permeability to Glu^-^ compared to Cl^-^. Group 4: Na^+^ substitution by Tris^+^ has no effect on V_rev_, indicating a loss of cation permeability. The V_rev_ shift in Tris^+^Glu^-^ vs. Tris^+^Cl^-^ was further strongly increased when the S^342^K mutation was combined either with D^352^K alone or with E^14^K and D^356^K together, an effect that could not be surpassed by mutating all three acidic residues to lysines or by additionally incorporating the S^339^A mutation.

To determine the possible role of the S342 gate (15) in the Na^+^ *versus* Cl^-^ selectivity of hP2X7R, we performed the above protocol on the serine-to-lysine substituted mutant hP2X7R^S342K^. The V_rev_ shift of hP2X7R^S342K^ in Na^+^Cl^-^ *versus* Tris^+^Cl^-^ was only -18 mV (compare Fig. 2 B and E), much smaller than the -48 mV shift observed for hP2X7R^wt^ (compare Fig. 2 A and D). The mean calculated permeability ratio P_Tris_:P_Na_ of 0.52 instead of 0.15 for the wt (*SI Appendix*, Table S1) indicates a reduced but still present preference for Na^+^ over the bulky Tris^+^ cation. This could mean that hP2X7R^S342K^ conducts Na^+^ worse or Tris^+^ better than hP2X7R^wt^ or both, with the degree of hydration relative to the pore size being a possible reason (28). Alternatively, the reduced pore size due to the S342K mutation may cause Tris^+^ to act as a “permeant-blocking ion “ (29) by tightly interacting with the pore and sterically hindering Na^+^ permeation, directly explaining the altered permeability ratio.

After additional substitution of Cl^-^ by Glu^-^ in the Tris^+^-based extracellular solution, V_rev_ remained virtually unchanged from -21.0 mV (Fig. 2B) to -21.6 mV (Fig. 2C), indicating that the reduced Na^+^ *versus* Tris^+^ preference was not associated with a significant Cl^-^ permeability of hP2X7R^S342K^.

### The hP2X7R^E14K,S342K,D352K^ triple mutant is an ATP^4-^-gated anion channel

As suggested from the cryo-EM structure, cations that have passed through the tri-S342 ring exit the central channel through cytoplasmic fenestrations (14). The MOLEonline program also predicts, with variable frequency and also depending on the homology mutant analyzed (almost 100% for hP2X7R^E14K^), lateral pathways from the large cavity below S342 lined by acidic residues such as E14 and D352 at the intracellular membrane interface of hP2X7R^wt^ (*SI Appendix*, Fig. S3). To determine the role of these conserved lateral acidic residues on ion selectivity, we mutated E14 and D352 to lysine together with the tri-S342 gate, yielding hP2X7R^E14K,S342K,D352K^. Replacing Na^+^ with Tris^+^ caused a statistically insignificant shift of V_rev_ of hP2X7R^E14K,S342K,352K^ towards positive values (compare Fig. 2C and F; for statistical analysis see Fig. 3). This suggests that the E^14^K,S^342^K,D^352^K mutation either eliminated the inherent cation selectivity of hP2X7R^wt^, or at least its preference for the small cation Na^+^ over the larger Tris^+^. Consistent with a complete loss of cation permeability, the additional substitution of extracellular Cl^-^ by the twice as large Glu^-^ was accompanied by a large positive V_rev_ shift of +39.9 mV (Fig. 2I), indicating a 4.8-fold greater Cl^-^ than Glu^-^ permeability (P_Glu_ : P_Cl_ of 0.21, mean 0.22 ± 0.05, see *SI Appendix* Table S1). It follows that lysine substitution of E14 and D352 (in addition to S342) in hP2X7R^E14K,S342K,352K^ results in selectivity for the small anion Cl^-^, in contrast to hP2X7R^wt^ and hP2X7R^S342K^, which are equally impermeable to Cl^-^ and Glu^-^. Both together, the insignificant V_rev_ shift due to Na^+^ by Tris^+^ substitution and the strong positive shift due to Cl^-^ by Glu^-^ substitution, are typical features of anion-selective ion channels such as the volume-regulated anion channel VRAC (30). We conclude that the hP2X7R^E14K,S342K,D352K^ mutant behaves like an ATP^4-^-gated anion channel.

### Acidic residues at the cytoplasmic exit of the TM2 channel control cation selectivity

To elucidate the individual contribution of the lysine substitutions of residues E14, S342, and D352 to the altered charge selectivity, we substituted each of these residues individually and in various combinations with lysines, including the non-conserved residue D356 and also S339, one helical turn above the gating residue S342 (see Fig. 1). Since S339K mutants were generally not functional (see also *SI Appendix*, Fig. S4), we had to make do with alanine mutants at this position. The reversal potentials were determined using the protocol described in Fig. 2. The data are summarized in Fig. 3A, sorted into 4 groups with comparative statistical analysis.

Group 1 shows the mean V_rev_ values of hP2X7R^wt^ in Na^+^Cl^-^ (white bar), Tris^+^Cl^-^ (red bar), and Tris^+^Glu^-^ (purple bar). Group 2 shows the V_rev_ values of hP2X7R mutants with single, double and triple lysine substitutions of E14, D352 or D356. Each single lysine substitution within this group consistently resulted in a less negative V_rev_ in Tris^+^Cl^-^ (red bars) compared to hP2X7R^wt^, indicating that each of the three mutations reduced the preference for Na^+^ relative to Tris^+^, however, Na^+^ was always preferred to Tris^+^ according to the still negative V_rev_ shift induced by the substitution of Na^+^ by Tris^+^. Replacing Tris^+^Cl^-^ with Tris^+^Glu^-^ did not significantly reduce the V_rev_ of the single mutants, indicating that the anion exclusion typical of hP2X7^wt^ is maintained (compare red and purple bars). However, when the E^14^K mutation was combined with D^352^K or/and D^356^K mutations, V_rev_ shifted statistically significantly to the positive in Tris^+^Glu^-^, i.e. when Cl^-^ was replaced by the organic anion Glu^-^ (compare height of purple *versus* red bars in Fig. 3A). This indicates that replacement of two or three acidic residues at the membrane-cytoplasmic interface of each subunit with lysines renders hP2X7R anion permeable.

Group 3 mutants, which address the contribution of TM2 residues S339 and S342 to ion selectivity, have less negative V_rev_ values in Tris^+^Cl^-^ than groups 2 and 1. The S^342^K mutation alone resulted in a less negative V_rev_ shift than any of the E^14^K, D^352^K, or D^356^K single mutations in group 2 when Na^+^ was replaced by Tris^+^ (heights of the red versus white bars). This indicates that S342 contributes significantly more to the small cation Na^+^ *versus* Tris^+^ selectivity of hP2X7R than either E^14^, D^352^, or D^356^ alone; this can best be explained by S342 acting mainly as a size selectivity filter, reducing the permeation of larger cations such as Tris^+^.

When the S^342^K mutation was combined with either the E^14^K or D^356^K mutation, an anion permeability was obtained, as evidenced by the statistically significant positive shift of V_rev_ upon Cl^-^ to Glu^-^ substitution. Since hP2X7R^S339K^ was not functional as an ATP^4-^-gated channel, we also examined the alanine mutant hP2X7R^S339A^. hP2X7R^S339A^ was functional but had no further effect on V_rev_ when combined with E^14^K and S^342^K (Fig. 3A). This suggests a negligible contribution of the hydroxyl side chain of S339 to the ion selectivity of hP2X7R.

The combination of the S^342^K and D^352^K mutations in hP2X7^S342K,D352K^ resulted in an even further shift to more positive V_rev_ values in Tris^+^Cl^-^ so that they were no longer different from the values in Na^+^Cl^-^ and were therefore assigned to group 4 (Fig. 3A). Furthermore, a particularly strong shift of V_rev_ to high positive voltages occurred here when Cl^-^ was replaced by Glu^-^ (compare the height of the purple bars between groups 3 and 4). The positive V_rev_ shift was further increased by including the E^14^K mutation in the triple mutant hP2X7^E14K,S342K,D352K^, whereas the triple mutant hP2X7^E14K,S342K,D356K^ was approximately equivalent to the double mutant hP2X7^S342K,D352K^. Taken together, this suggests that D352 is more important for ion selectivity than E14 or D356, both of which had to be simultaneously mutated to lysines to have the same effect as D^352^K when combined with S^342^K. The lack of effect of the Na^+^ to Tris^+^ substitution on V_rev_ combined with the simultaneous strong shift of V_rev_ to positive potentials in Tris^+^Glu^-^ indicates that all group 4 hP2X7R mutants are purely selective for the small Cl^-^ anion.

A complete shift from cation to anion permeability was also induced by mutation of D352 or/and D356 in the E^14^K,S^342^K background. Again, the additional alanine mutation of S339, which is one helical turn upstream of S342 and, like S342, is located in the narrowest region of the closed trihelical TM2 channel (Fig. 1), had no additional effect on the positive V_rev_ shift induced by the Cl^-^ to Glu^-^ substitution (Fig. 3). This supports our previous single-channel data that S339 does not act as a selectivity filter (15), and is also consistent with the cryo-EM structure and our derived homology models showing that the open pore diameter is significantly larger above and below the triple S342 ring, except for the constriction at K17, which can be ruled out as an exit site (see *SI Appendix*, Fig. S2). Unfortunately, the non-functional state of hP2X7R^S339K^ did not allow a more comprehensive analysis. The alanine and lysine mutants of Y343, next to S342 in the ion channel, were also not functionally expressed. The permeability ratios reflecting the V_rev_ shifts of all hP2X7R mutants examined are summarized with statistical analysis in *SI Appendix*, Table S1.

The ATP^4-^-induced conductances (G_at Vrev_) of all hP2X7R constructs are shown in the right part of Fig. 3. The ATP^4-^-induced conductances differed greatly between the different hP2X7R mutants. In contrast, the biochemically visualized plasma membrane expression of the hP2X7R mutants examined varied muss less to be considered a major cause of the conductance variations. Therefore, functional reasons must be responsible for the variable conductances.

### Mutation of acidic residues to lysine at the cytoplasmic interface causes strong inward rectification

The I-V curves in Fig. 2 A,D,G show a nearly linear current-voltage relationship for ATP^4-^-induced currents mediated by hP2X7R^wt^, indicating little or no rectification. This is consistent with previous measurements in both whole-cell mode (5, 8) and single-channel mode (10, 27, 31). In contrast, the I-V curves for hP2X7R^S342K^ (Fig. 2 H) show strong outward rectification, which was reversed to inward rectification by additional lysine substitutions of E14 and D352 in the triple mutant hP2X7R^E14K,S342K,D352K^ (Fig. 2 I). To systematically examine the correlation, we quantified the rectification index for hP2X7R^wt^ and all its mutants as the ratio of the slope conductance at negative and positive potentials in all three extracellular solutions (see Materials and Methods), as shown as examples for hP2X7R^S342K^ and hP2X7R^E14K,S342K,D352K^ (*SI Appendix*, Fig. S5A and Fig. S5B, respectively). Statistics are presented as bar graphs (*SI Appendix*, Fig. S5C). In almost all constructs summarized as group 2 in Fig. 3, outward rectification increased significantly when Na^+^ was replaced by Tris^+^. This can be explained by a decrease in the inward current, which is now carried by the larger, less permeable Tris^+^ ion instead of Na^+^, providing additional evidence for small cation selectivity. The strong inward rectification of the hP2X7R^E14K,S342K,D352K^ mutant in Tris^+^Glu^-^ (reflected by the absence of outward current at positive potentials up to +40 mV) can be explained by the absence of available charge carriers: endogenous intracellular Cl^-^ can move out at negative potentials, but Glu^-^ cannot enter because of its larger size.

### Effect of lysine mutations on pore diameter and shape by homology modeling and structural visualization

A compilation of SWISS homology**-**modeled pore sections shows that the isosceles triangular arrangement of residues is preserved in all 17 electrophysiologically characterized hP2X7R mutants, even with multiple lysine substitutions. The open states of wt and five of these mutants are exemplarily shown in *SI Appendix*, Fig. S4, in top (A-F) and bottom views (G-L) together with the modeled pores (M-R). The S342K mutation (*SI Appendix*, Fig. S4, compare A-C with D and E), but not the non-functional S339K mutation (*SI Appendix*, Fig. S4, compare A with F), reduced the visible pore diameter at the level of residue 342 by about half (from 6.0 Å to 3.0-3.3 Å). In contrast, pore diameter at residue 342 was unaffected by multiple acid-to-lysine mutations in the acidic triangle (*SI Appendix*, Fig. S4, compare A-C).

## Discussion

Based on the structure of the truncated homotrimeric zebrafish P2X4R (32, 33), the narrowest part of the ATP-gated ion-conducting pathway is formed by the trihelical TM2 bundle, as confirmed by functional experiments (34) and also by the cryo-EM structure of rP2X7R (14). In hP2X7R, we found the tri-S342 ring in TM2 to be involved in gating and permeation, acting as a size selectivity filter (15). Our new results show that the S342K mutation only affects the relative selectivity for Na^+^ *versus* Tris^+^, which have effective diameters of 1.9 Å versus 6.5 Å, respectively, but does not itself induce anion permeability. This is in contrast to the reported importance of the equivalent residue Thr339 in rat P2X2R, which when mutated to lysine changed permeation from 10-fold cation selective to anion preferential (35).

### Acidic isosceles triangle: a large-diameter, high field electrostatical cation selectivity filter for small inorganic and small organic cations

In our previous SCAM experiments (15), we observed that the membrane-impermeant anionic cysteine-reactive 5.8 Å reagent MTSES^-^ reacted with S342C when applied externally, but not when applied to the cytosolic side. In contrast, the cationic 5.8 Å reagent MTSET^+^ reacted with S342C from both sides of the membrane. A logical explanation for this difference is the presence of a true cation-selective filter downstream of S342 that is impenetrable to the MTSES^-^ anion from the cytosolic side. Based on the recognition of the closely spaced acidic residues E14, D352 and D356 deep in the membrane at the cytoplasmic interface, we were able to gradually shift the permeation properties of hP2X7R from cation to anion selectivity by stepwise mutation of the acidic residues to lysines. The importance of basic residues as anion selectivity filters is well known for instance for TMEM16A Ca^2+^-activated Cl^-^ channels (36).

Considering the ability of the 5.8 Å diameter MTSET^+^ to react “backwards” with S^342^C when applied cytosolically (15), we can retrospectively conclude that the acidic cation selectivity filter at the channel exit must also have a diameter of at least 5.8 Å. Calculation of the mean distances by PyMol between the carboxyl groups of E14 and D352 and E14 and D356 of the homology-modeled hP2X7R mutants gave 9.6 ± 1.5 Å and 5.4 ± 0.5 Å (± SD), respectively, and 7.4 ± 2.5 (± SD) overall, which is in the range consistent with anterograde Tris^+^ and retrograde MTSET^+^ permeation. We propose that the high density of three negative charges per lateral pore at the interface of two subunits is necessary to generate an electrostatic field strong enough to force selectivity even for solvated cations. In other words, the high charge density serves to compensate for the large pore radius. This view is supported by our observation that all three acidic residues (together with the size filter at S342) are necessary for pure cation selectivity and that this selectivity gradually disappears with each acidic residue mutated to a lysine.

### Tri-S342 ring in membrane center: primarily a size filter for cations

Since full anion selectivity required the additional S342K mutation in the E^14^K/D^352^K/D^356^K background, and also for electrostatic reasons and the reduced modeled pore size of the S342K mutants to 3.3 Å (*SI Appendix*, Fig. S3), close to the 3.6 Å ionic diameter of Cl^-^, we considered that the S342K mutation alone would convert the hP2X7R from Na^+^ to Cl^-^ permeability. However, no detectable anion permeability was seen with the S342K mutation alone. How the S342K mutation, together with the lysine substitutions in the acidic triangle, contributes to Cl^-^ selectivity remains unclear at this time.

### Role of acidic residues in the TM2 of P2XRs in relation of literature

The basic data of this work are graphically summarized in Fig. 4. Based on the insight that the arrangement of acid residues in the cytoplasmic interface may serve as a selectivity filter, we were able to gradually change the permeation characteristics from selectivity for small cations (hP2X7R^wt^) to selectivity for small anions (e.g., hP2X7R^E14K,S342K,D352K^). The virtual impermeability to anions provides additional support for the view that the proposed dilation of the P2X7R ion channel pore, reportedly associated with the ion channel becoming non-selective also for large organic cations (5, 37) and anions (Browne et al. 2013), is an artifact. Our new data re-emphasize the view that the hP2X7R maintains a stable conductance for cations with a minimum cross-sectional diameter < 8.5 Å (10, 15), but is per se impermeable to anions (8) (27).

**Fig. 4.**
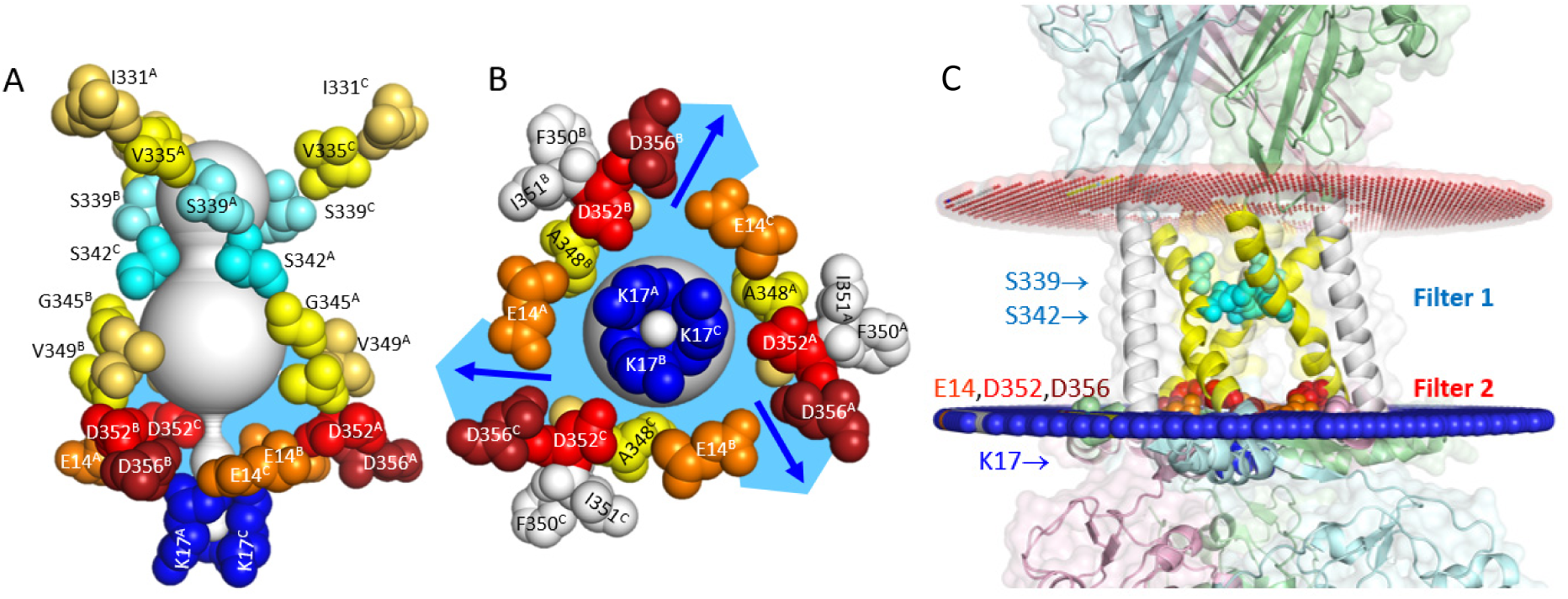
Cation selectivity filter at the cytoplasmic interface of hP2X7R^wt^. Lateral (A) and bottom (B) views of the open hP2X7R^wt^ with labeled residues lining the open pore or located more peripherally (F350, I351, colored gray). The pore was modeled using MOLEonline. From the frequent detection of single lateral pores in the open hP2X7R channel by the MOLEonline software (see *SI Appendix*, Fig. S3) and consistent with the data shown here, we conclude that the fluid in the cavity below the tri-S342 ring (sky-blue) and the lateral pores is continuum, with access controlled by the gate at S342 (selectivity filter 1). C, Location of filters 1 and 2 relative to phospholipid head groups. Our electrophysiological data indicate that cations exit through three exits (indicated by blue arrows), one between each subunit interface and each flanked by E14 of one subunit and D352/D356 of the adjacent subunit (size selectivity filter 2). The membrane boundaries of the SWISS homology modeled open hP2X7R^wt^ were determined using the “Orientations of Proteins (OPM) in Membranes” server (https://opm.phar.umich.edu/) with the conditions “Mammalian plasma membrane”, “Allow curvature yes”, “Topology N-term in” (24, 59). The red and blue pseudo-atoms were automatically added by the OPM server to mark the hydrophobic boundaries of the lipid bilayer. The downloaded PDF file was visualized with PyMol. Note the proximity of the cation selectivity filter to the phospholipid headgroups and the apparent lack of coverage of the transmembrane channel by the main and side chains of flanking amino acid residues, both of which may be related to the effect of lipids on ion permeation (13) (channel portion in (A) uncovered by amino acid main and side chains).

Here, we show that in addition to S342 of TM2, ions are selected by acidic amino acid residues (E14, D352 and D356) along the lateral pores of the cytoplasmic part of the hP2X7R protein. Lysine mutations at S342 and at the critical lateral pore residues produced ion channels with currents independent of extracellular cations, indicating a loss of cation permeability. Instead, currents using the “new” selectivity filters at K342 and K352 (or additionally at K14 and K356) are dependent on the size of the extracellular anion, i.e. the filters allow Cl^-^ (effective diameter 3.6 Å) to permeate more easily than the bulkier Glu^-^ (effective diameter 5.5 Å). The lateral pores are part of the cytoplasmic cap (14) and allow ions to exit the large cavity below S342 in the cytoplasmic half of the membrane through the size-selective filter into the cytoplasm. However, a complete reversal of charge selectivity from cationic to anionic required mutations of S342 and at least D352 to lysine, suggesting that the cation selectivity of hP2X7R is a product of the combination of the two filters.

Our results indicate that D352 of P2X7R or equivalent positions of other P2XRs are critical for P2XR ion permeation. It has been suggested previously that ion flow through P2Xs into the cytoplasm does not occur along the central axis of the cytoplasmic domain, but through lateral fenestrations (14, 38), but which amino acid residues line these pores remained unresolved. Several previous reports support our view of the important role of D352 in the P2X7R ion permeation pathway. Single channel currents of rat P2X2R were abolished when residue D349, corresponding to D352 in hP2X7R (and D361 in the 12 N-terminal residues longer hP2X2R, *SI Appendix*, Fig. S6), was replaced by neutral or positively charged residues (39). rP2X2R^D349C^ mediated only small currents after activation by ATP (40) and was blocked by MTSEA^+^ (41) or Cd^2+^ (42). These results indicated early on that this position is critical for ion permeation. Furthermore, hP2X7R^D352C^ mediated only 10% of the hP2X7R^wt^ conductance (15). Substitution of T348 and D352 with basic residues in the channel-lining TM2 domain of the rP2X7R simultaneously increased the permeability of the normally cationic channel for Cl^-^ and an acidic fluorescent dye with an effective diameter of >10 Å (43).

Residues equivalent to E14 of hP2X7R were previously identified as critical for the cation selectivity of rat P2X2 (E17) and the additional anion permeability of mouse P2X5 (K17), as well as for the access to the transmembrane pore through lateral fenestrations (20). The importance of E14 for the Ca^2+^ permeability of rP2X7R has already been shown (44). In an alternatively spliced version of P2X7 (P2X7k (45)), the acidic E14 is replaced by the neutral asparagine residue. This leads to a shift of the reversal potential to more positive potentials in NMDG^+^ containing extracellular solution. The fractional P2X7k-mediated Ca^2+^ current is reduced compared to hP2X7^wt^ (44). These results previously suggested that the N-terminal E14 residue is important for the permeation properties of hP2X7R.

The involvement of lateral fenestrations in the extracellular domain in the ion selection process has been proposed for the P2X4R (46-48). Although ions may use a similar pathway in P2X7R-mediated currents, the extracellular pores appear to be irrelevant for the cation selectivity of P2X7R, since the combined mutation at the two filter sites, S342 and acidic triangle, is sufficient to completely alter the charge selectivity of hP2X7R ions.

All these previous findings, together with our results shown here, support our view that ion selection of P2X7Rs occurs in a two-step fashion, with (i) a selectivity filter located in the middle of the TM2 domain and (ii) acidic residues lining lateral fenestrations controlling cation flow from the bulky cavity in the cytoplasmic half of the membrane into the cytoplasm (see graphical summary in Fig. 4). The conservation of D352 across the P2XR family (*SI Appendix*, Fig. S6; see also a comprehensive interspecies P2X sequence alignment in (32)) suggests that the cation selectivity of P2XR generally involves the mechanism described here.

## Materials and Methods

### Reagents

Unless otherwise stated, standard chemicals were purchased from Merck/Sigma Aldrich (Darmstadt, Germany). Na_2_ATP was purchased from Roche (Mannheim, Germany). Molecular biology reagents were purchased from New England Biolabs (Schwalbach, Germany) with the exception of the anti-reverse cap analog m7,3’-OGpppG (product NU-855), which was purchased from Jena Bioscience, Germany.

### Homology modeling of the hP2X7 homotrimer and point mutants in the apo-closed and open states

According to a ClustalW alignment (49), the primary sequence identity between hP2X7 (Q99572) and rP2X7 (Q64663) is 80.3%. Using Swiss-Modeler (https://swissmodel.expasy.org) and the cryo-EM structures of rat P2X7R (rP2X7R) in the apo-closed state (PDB 6U9V) and in the ATP-bound open state (PDB 6U9W) as templates (14), we generated homology models of hP2X7R^wt^ and mutants and visualized them using PyMol (Schrödinger LLC. The PyMOL molecular graphics system, version 1.3r1. Portland. 2010 Oregon: Schrödinger, LLC.). To calculate pores and tunnels in the homology-modeled hP2X7R structures, we used MOLEonline (25) and visualized them using the built-in LiteMole viewer or after downloading the results in PyMol format using PyMol software.

### Generation of hP2X7R mutants

The plasmid encoding the wild-type human P2X7R (hP2X7R) subunit (P2RX7_HUMAN, UniProtKB Q99572) in our oocyte expression vector pNKS2 (50) was the same as in our previous studies (9, 10, 15, 51). Point mutants were designed using Vector NTI software (InforMax v. 4.0) and generated using the QuickChange method for site-directed mutagenesis (52). The primers used are listed in *SI Appendix*, Table S2. Mutants were screened by restriction pattern analysis and verified by commercial DNA sequencing (MWG, Ebersberg, Germany) and sequence alignment. Capped and polyadenylated cRNAs were synthesized from XhoI linearized DNA templates as described previously (53).

### Expression of hP2X7R^wt^ and its mutants in *X. laevis* oocytes

The maintenance of the frogs and the surgical removal of parts of their ovaries were approved by the local animal care committee (ref no. 42502-2-1493 MLU) according to EC Directive 86/609/EEC for animal experiments. After defolliculation with collagenase NB 4G (Nordmark Pharma GmbH, Uetersen, Germany), Dumont stage V and VI oocytes were injected with 46 nl of cRNA containing 1 ng or 50 ng of wild-type or mutant hP2X7R, respectively. Injected oocytes were maintained at 19°C in oocyte Ringer’s solution (ORi: 100 mM NaCl, 1 mM KCl, 1 mM MgCl_2_, 1 mM CaCl_2_, 10 mM HEPES-NaOH, pH 7.4) supplemented with penicillin (100 U/ml) and streptomycin (100 μg/ml) until used 1 - 3 days later.

### Two-electrode voltage-clamp (TEVC) recordings from *X. laevis* oocytes

Currents were recorded at room temperature (∼22 °C) using an OC-725C oocyte clamp amplifier (Warner Instruments, Hamden, USA), filtered at 100 Hz, and sampled at 500 Hz as previously described (19, 27, 53). The reference electrodes were connected to the bath via 3 M KCl agar bridges. The remaining diffusion potentials were measured according to a previously published method (54). In the Tris^+^-Glu^-^-based extracellular solution, this potential was −8 mV and was corrected accordingly. The diffusion potential for Cl^-^-based bath solutions was negligibly small (< 0.5 mV).

While superfused with ORi, individual oocytes were impaled with 3 M KCl-filled glass microelectrodes with resistances of 1.2–1.4 MΩ. hP2X7R-dependent currents were measured in nominally Ca^2+^- and Mg^2+^-free bath solutions consisting of either 100 mM NaCl, 100 mM TrisCl or 100 mM Tris-glutamate supplemented with 0.1 mM flufenamic acid, 1 mM EGTA and 5 mM HEPES-NaOH, pH 7.4. Flufenamic acid was used to block the conductance caused by removal of external divalent cations (55-57). The hP2X7R-mediated inward currents were elicited by switching for 5 s to the same bath solution containing an additional 0.1 mM free ATP (ATP^4-^). The interval between ATP applications was 2 min. Each of the three different bath solutions was applied once to the same oocyte in random order. Switching between the different bath solutions was accomplished within < 1 s by a set of computer-controlled solenoid valves combined with a modified U-tube technique (8).

Ramp currents were measured during 500 ms long voltage ramps applied every 1 s between −80 mV and +40 mV. The holding potential was maintained at −40 mV between ramps. ATP-induced ramp currents were calculated as the difference between the ramp currents before and during ATP application.

### Data analysis and presentation

Data were stored and analyzed on a personal computer using software developed in our department (Superpatch 2000, SP-Analyzer by T. Böhm, Julius-Bernstein-Institute of Physiology, Halle, Germany). SigmaPlot (Systat Software) was used for fitting, statistical analysis, and data presentation. Statistical data are expressed as mean ± SEM and analyzed by one-way ANOVA. Bonferroni’s t-test for multiple comparisons was used to test the statistical significance of differences between means. Statistical significance was set at a p-value of <0.05.

The permeability ratios (P_X_/P_Y_) were calculated from the changes in reversal potential by the Goldman equation (26):

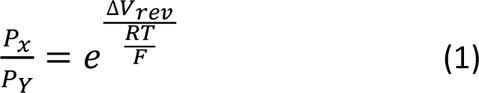

where x and y are cations (Na^+^ or Tris^+^) or anions (Cl^-^ or Glu^-^) respectively, and ΔV_rev_ is the reversal potential difference measured in extracellular Na^+^Cl *versus* Tris^+^Cl^-^ or Tris^+^Cl^-^ *versus* Tris^+^-Glu^-^-based solutions.

The degree of rectification of the I-V curves, also known as the rectification index, was calculated as the ratio of the outward to inward conductances, G_outward_/G_inward_, as described previously (58). The G_outward_ and G_inward_ values were determined from the slopes of the I-V curves in the voltage ranges from −75 to −40 mV and from +20 to +40 mV (*SI Appendix*, Fig. S5).

## Acknowledgments

F.M. and G.S. thank the Deutsche Forschungsgemeinschaft for financial support through grants MA 1581/15-2, SCHM 536/9-2, and SCHM 536/12-1, respectively.

## Institutional Review Board Statement

The procedures for maintenance of the frogs and their ovariectomy were approved by the local animal welfare committees (Halle, Germany, reference no. Az. 203.42502-2-1493 MLU, and Düsseldorf, Germany, reference no. 8.87-51.05.20.10.131 for experiments performed in Halle and Aachen, respectively) in compliance with EC Directive 86/609/EEC for animal experiments.

## Figures

**SI Appendix, Fig. S1.**
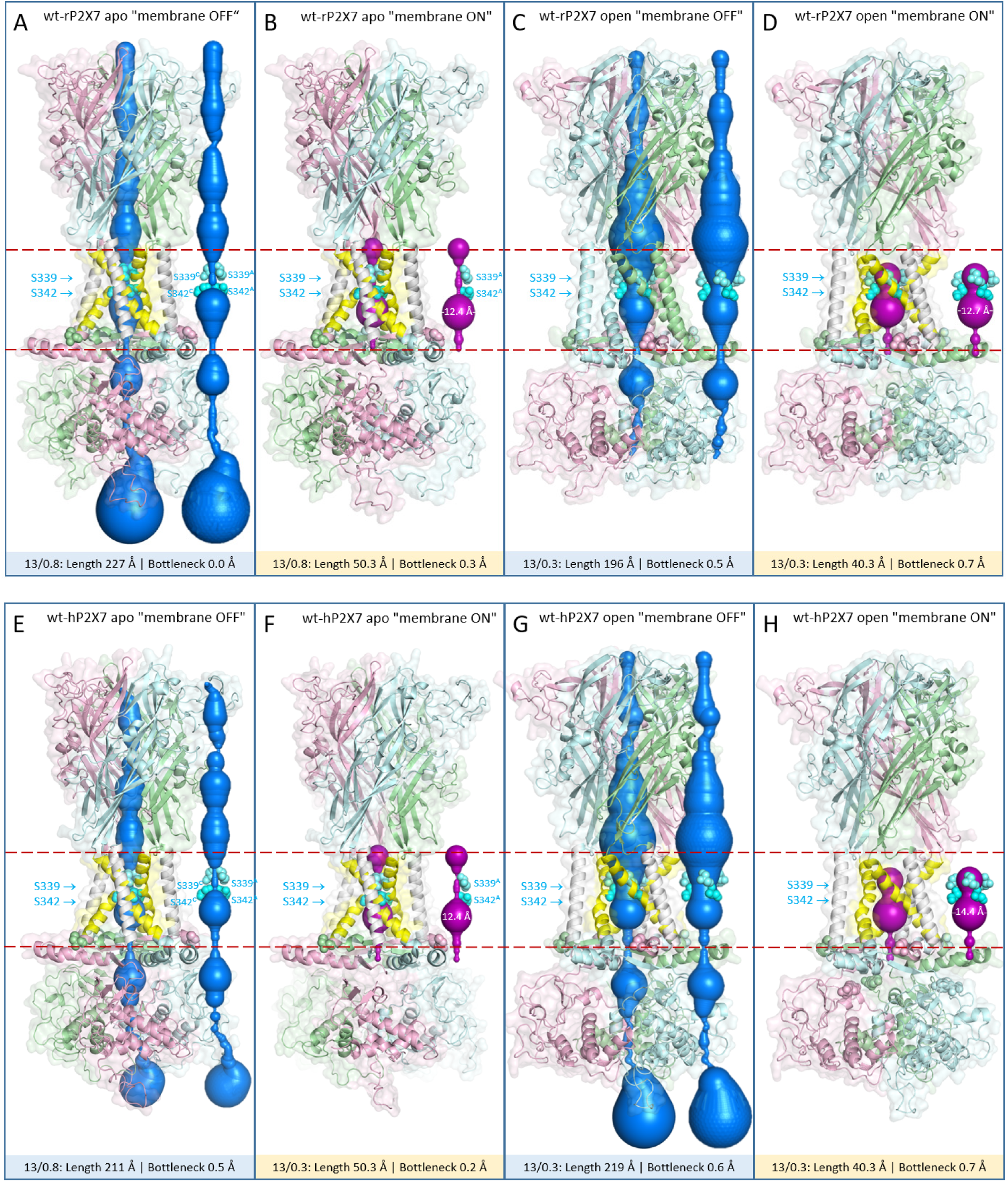
Pore structures of rP2X7R and hP2X7R calculated with the MOLEonline program without and with consideration of the position of the plasma membrane. Shown are side views based on the cryo-EM structure of rP2X7R (A-D) and the derived SWISS homology model of hP2X7R (E-H). The calculated pore structures are also shown without the surrounding protein to visualize details of the bottlenecks. When one or two serine residues are omitted for better visibility in A,B and E,F, the serine residues shown are labeled with their residue numbers and the corresponding polypeptide chain as superscript. The predicted outer and cytoplasmic borders of the plasma membrane are indicated by the dashed red lines. The figure was generated with PyMol.

**SI Appendix, Fig. S2.**
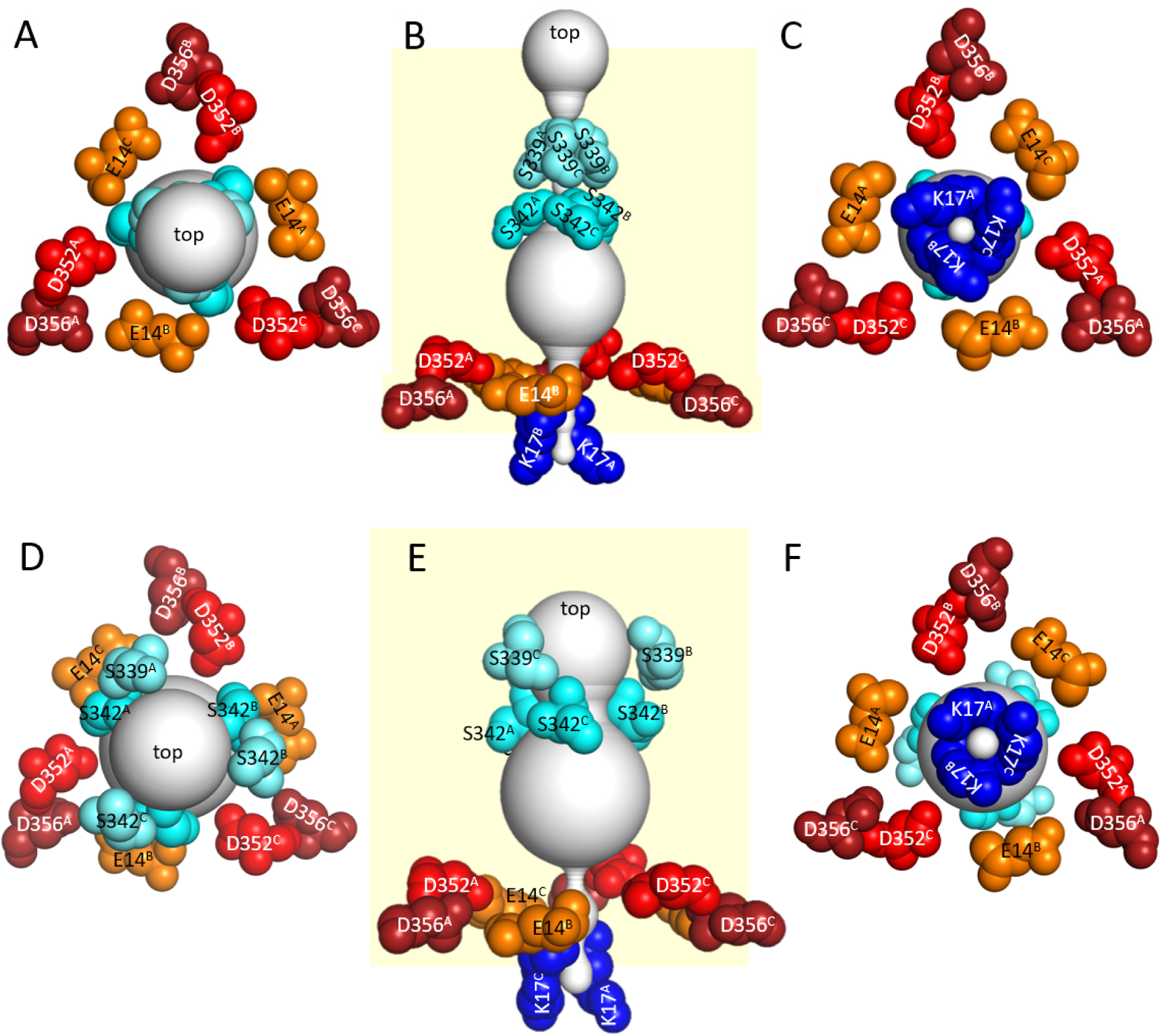
Location of acidic and basic residues at the cytoplasmic interface of hP2X7R^wt^ relative to the channel pore. Shown are top (A,D), side (B,E) and bottom views of the closed (A-C) and open (D-F) pores (in light gray) of hP2X7R as predicted by the MOLEonline program in the “membrane ON” configuration. The top views show the arrangement of the nine acidic residues, three from each subunit, in an isosceles triangular structure. The membrane position as predicted by the OPM database is highlighted in light yellow (B,E). In both the closed and open states, the single exit to the cytoplasm predicted by MOLEonline, formed by three circularly arranged K17 residues, is far too narrow to allow ions to exit from the central vestibule above. The figure was generated using PyMol.

**SI Appendix, Fig. 3.**
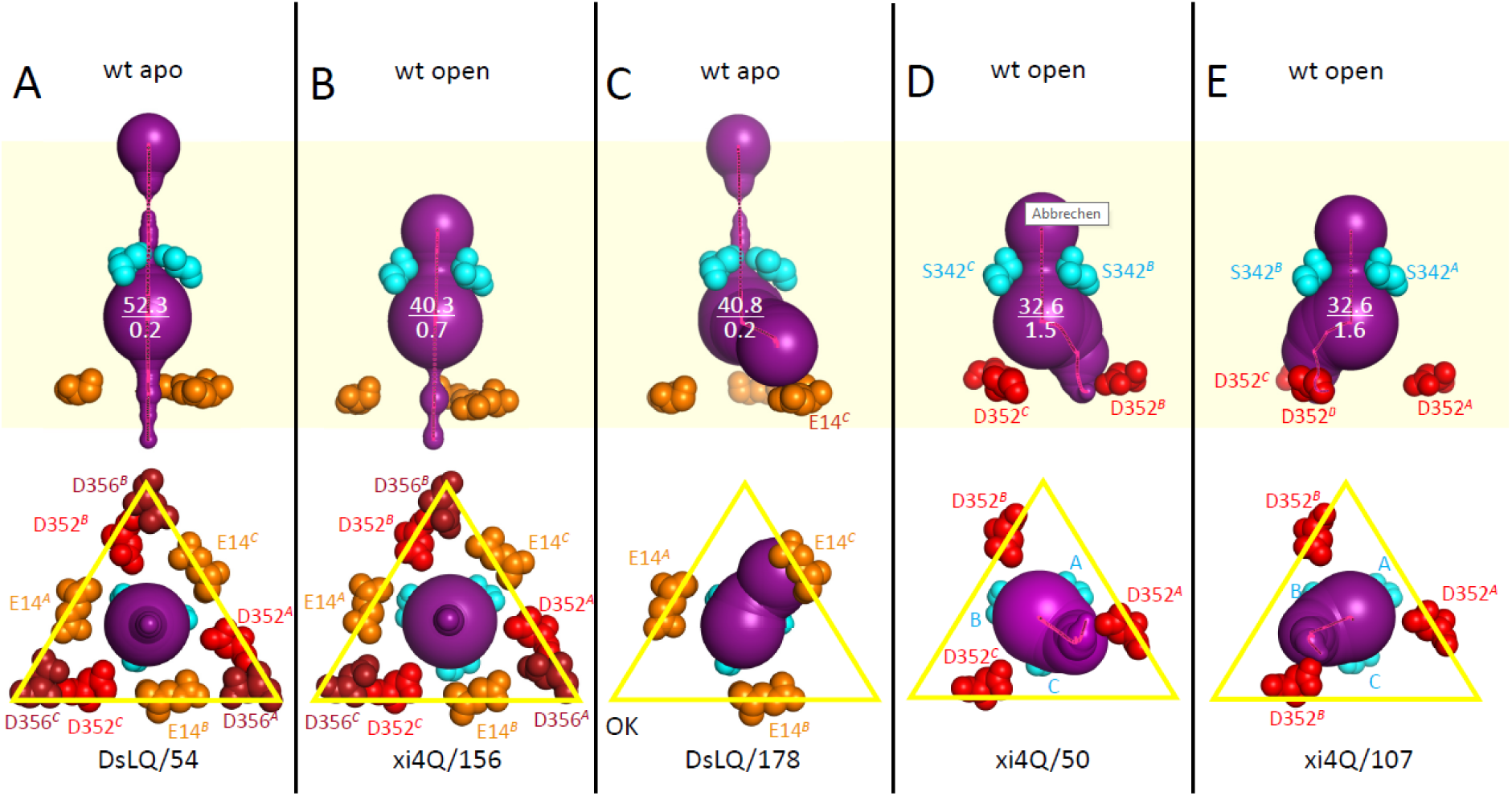
Lateral pores are randomly detected in the homology-modeled hP2X7R by MOLEonline. Closed and open hP2X7R^wt^ structures were repeatedly analyzed with identical or varying settings of the MOLEonline software in the “Membrane ON” configuration. A,B represent the prevailing results in the side and bottom views already shown in Fig. 1. The side view shows only the positions of the E14 residues, while the bottom view shows all nine acidic residues within the isosceles triangle. C-E show examples obtained with identical MOLEonline settings that, in addition to the consistently modeled large vestibule, have a modeled lateral pathway flanked by one of the critical acidic residues, E14^B^ (C), D352^A^ (D), and D352^B^ (E). Serine 342 residues are shown in cyan. The numbers at the bottom are part of the bookmarks under which the specified data can be reopened. The lateral pores discovered in C-E do not increase the diameter of the large cavity below S342. This suggests that the lateral pores only branch below the equator of the cavity. The figure was generated using PyMol. Repetitive MOLEonline calculations (60 each) with identical default settings revealed lateral paths at different frequencies depending on the hP2X7R variants studied: about 15% for hP2X7R^wt^, 38% for hP2X7R^E14K^, 33% for hP2X7R^S342K^, 100% for hP2X7R^D352K^, and 12% for hP2X7R^D356K^.

**SI Appendix, Fig. S4.**
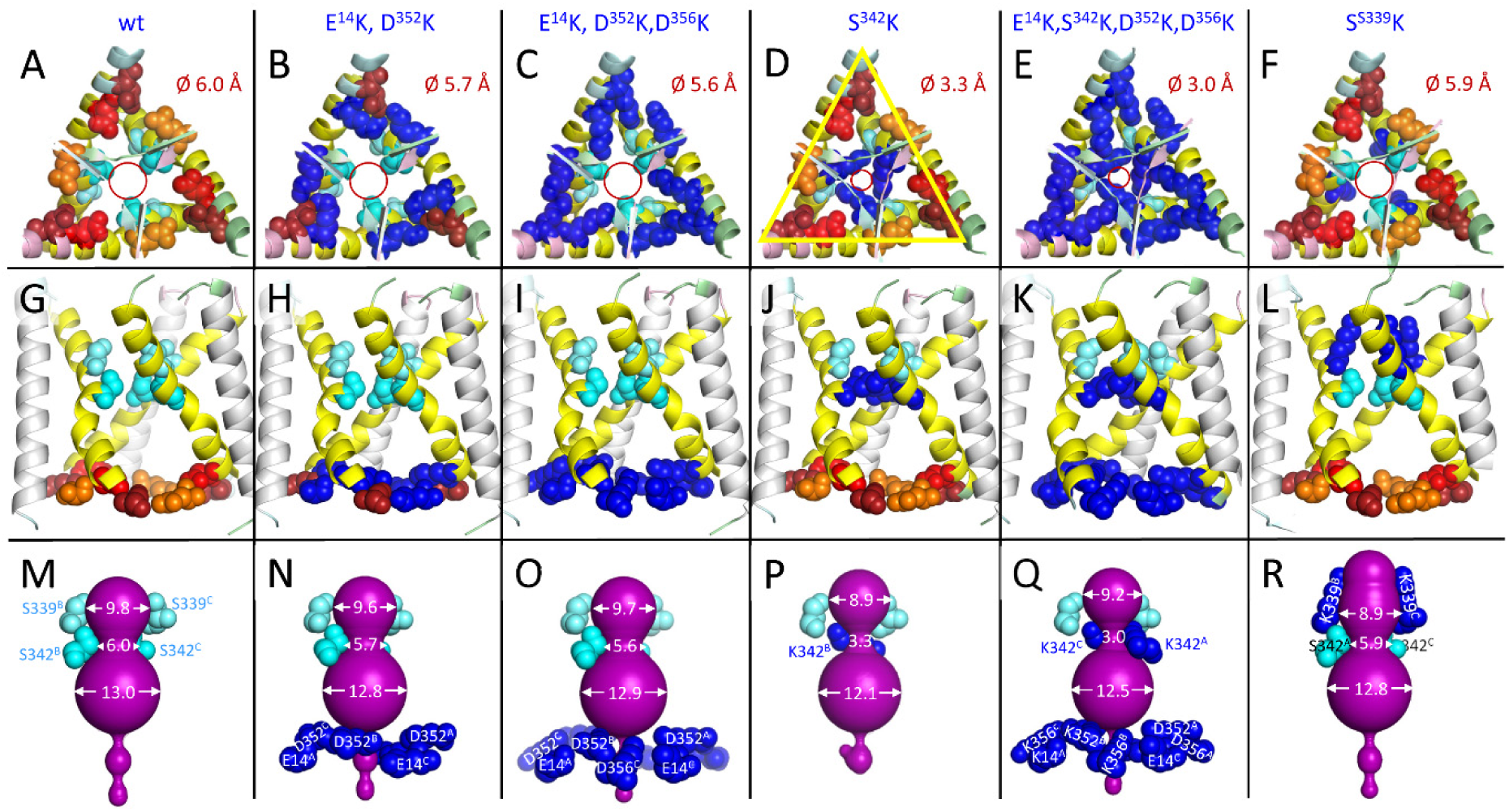
Pore size at the level of gating residue S342 is affected by lysine substitution of S342, but not by lysine substitution of acidic residues in the isosceles triangle at the cytoplasmic interface. A-F show the top and G-L the corresponding side views of the membrane-embedded region of the indicated SWISS homology-modeled open hP2X7R constructs. M-R show the corresponding pore structure (in purple) as predicted by MOLEonline. The indicated diameters (in Å) of the upper and lower vestibules and the pores at S342 were graphically derived from the MOLEonline-predicted channel profiles. Orange, red and firebrick colored residues represent E14, D352 and D356, blue colored residues represent lysines. All mutants are functional except S339K (far right row), which is not functional when mutated alone or in combination. The highly distorted shape of the upper vestibule (seen in R) may prevent extracellular cations from entering the central pore, but this was not further investigated and is therefore speculative.

**SI Appendix, Fig. S5.**
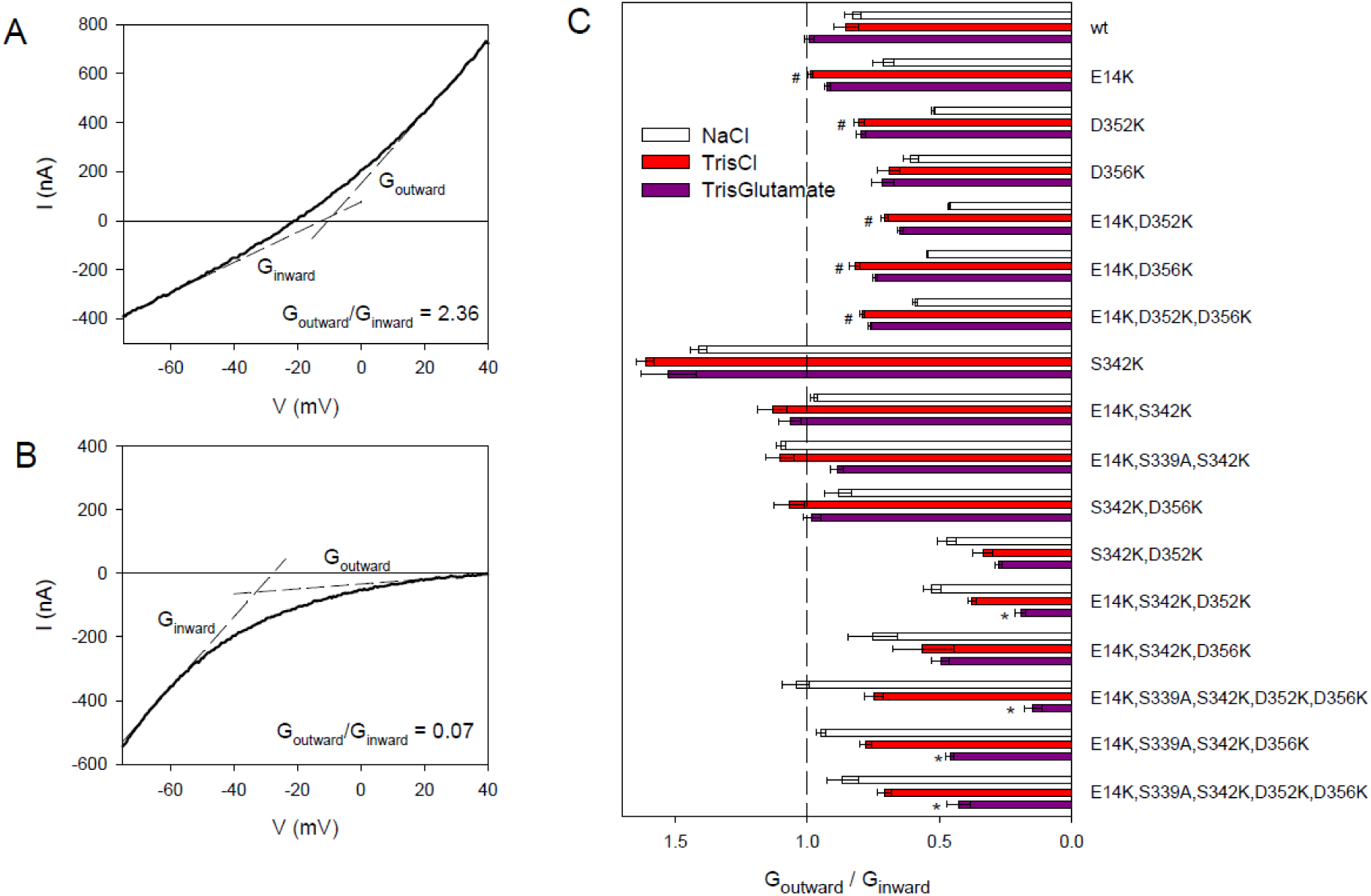
Rectification of hP2X7R^wt^ and mutants. A and B, examples of (A) outward rectification (hP2X7^S342K^ in Tris^+^Cl^-^) and (B) inward rectification (hP2X7R^E14K,S342K,D352K^ in Tris^+^Glu^-^). The rectification indices are shown in the figures. C, Statistics of rectification index G_outward_/G_inward_ for the hP2X7R constructs studied. Symbol # indicates a significant change in rectification when Na^+^Cl^-^ was replaced by Tris^+^Cl^-^. Symbol * indicates a significant decrease in rectification index when Tris^+^Cl^-^ was replaced by Tris^+^Glu^-^. Bars are mean ± SEM of 6-7 oocytes.

**SI Appendix, Fig. S6.**
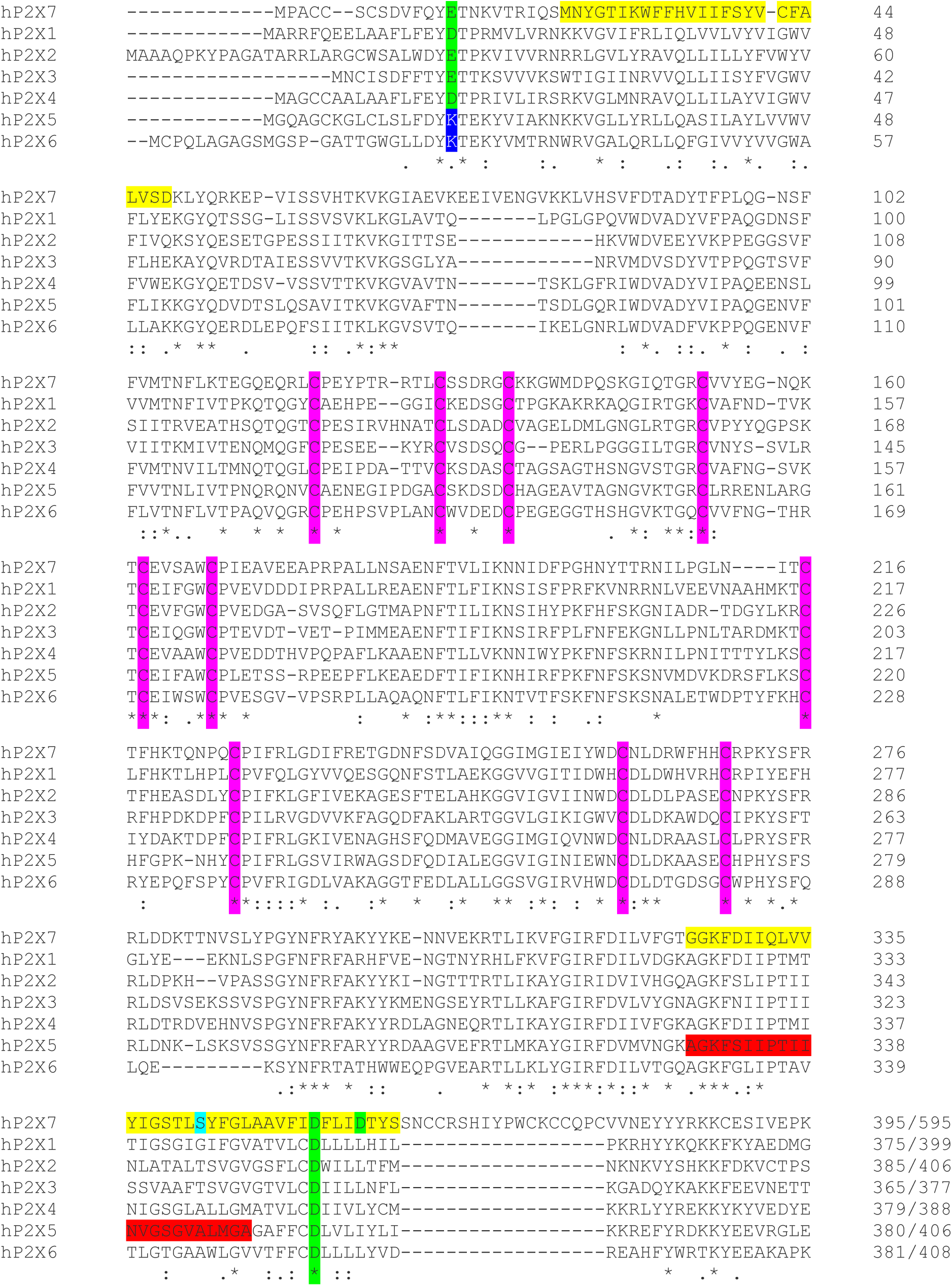
Protein sequence alignment of the seven hP2X isoforms. A protein blast search was performed using the 595-residue long hP2X7A subunit as the query sequence in the UNiProKB/Swiss-Prot database. Since the C-terminal endodomains of the different hP2X subunits show minimal homology, the alignment was terminated at position 395 of the 595 amino acid long hP2X7 protein chain. The conserved ectodomain cysteine residues are highlighted in magenta. In the hP2X7 sequence, residue S342 is highlighted in turquoise as the gate and acidic residues E14, D352 and D356 are highlighted in green as ion selectivity candidates. Note that E14 of hP2X7 is conserved in all other P2X subunits except hP2X5 and hP2X6, which have a lysine (K, highlighted in blue) at the corresponding position. The N-glycosylation sequons (NXS/T) are highlighted in gray. The UniProt/SwissProt accession numbers are P51575.1 (hP2X1); Q9UBL9.1 (hP2X2); P56373.2 (hP2X3); Q99571.2 (hP2X4); Q93086.4 (hP2X5); O15547.2 (hP2X6); Q99572.4 (hP2X7). Since the UniProt/SwissProt only contains the prevailing, but non-functional splice variant, which is missing 22 codons in the pre-TM2 and TM2 regions [1], the full-length sequence was completed with the missing residues (highlighted in red).

---

**SI Appendix, Table S1.** Tris^+^/Na^+^ and Glu^-^/Cl^-^ permeability ratios of wt and mutant hP2X7Rs. Permeability ratios were calculated using the Goldman-Hodgkin-Katz equation (Hille 1971) from V_rev_ values determined from I-V curves as shown in Fig. 2. Data are mean ± SEM of measurements on N different oocytes. Calculations are based on the mean reversal potentials shown in Fig. 3. The hP2X7R constructs are divided into the same groups as in Fig. 3 and numbered accordingly. Blank fields indicate that the corresponding hP2X7R constructs did not show significant V_rev_ changes after substitution of Na^+^ or Cl^-^ by Tris^+^ and Glu^-^, respectively, due to negligible cation or anion permeability. Since the derived permeability indices are not meaningful, they have been omitted.

| hP2X7R construct | Group in Fig. 3 | Tris<sup>+</sup>/Na<sup>+</sup> | SEM | N | Glu<sup>-</sup>/Cl<sup>-</sup> | SEM | N |
| --- | --- | --- | --- | --- | --- | --- | --- |
| wt | 1 | 0.15 | 0.01 | 6 |  |  |  |
| E<sup>14</sup>K | 2 | 0.29 | 0.03 | 6 |  |  |  |
| D<sup>352</sup>K | 2 | 0.35 | 0.03 | 6 |  |  |  |
| D<sup>356</sup>K | 2 | 0.27 | 0.01 | 6 |  |  |  |
| E<sup>14</sup>K,D<sup>352</sup>K | 2 | 0.41 | 0.04 | 6 | 0.59 | 0.05 | 6 |
| E<sup>14</sup>K,D<sup>356</sup>K | 2 | 0.40 | 0.04 | 6 | 0.53 | 0.05 | 6 |
| E<sup>14</sup>K,D<sup>352</sup>K,D<sup>356</sup>K | 2 | 0.29 | 0.02 | 6 | 0.68 | 0.04 | 6 |
| S<sup>342</sup>K | 3 | 0.52 | 0.04 | 11 |  |  |  |
| E<sup>14</sup>K,S<sup>342</sup>K | 3 | 0.61 | 0.08 | 6 | 0.57 | 0.04 | 6 |
| E<sup>14</sup>K,S<sup>339</sup>A,S<sup>342</sup>K | 3 | 0.63 | 0.04 | 6 | 0.58 | 0.02 | 6 |
| S<sup>342</sup>K,D<sup>356</sup>K | 3 | 0.57 | 0.03 | 7 | 0.67 | 0.08 | 7 |
| S<sup>342</sup>K,D<sup>352</sup>K | 4 |  |  |  | 0.23 | 0.04 | 5 |
| E<sup>14</sup>K,S<sup>342</sup>K,D<sup>352</sup>K | 4 |  |  |  | 0.22 | 0.05 | 6 |
| E<sup>14</sup>K,S<sup>342</sup>K,D<sup>356</sup>K | 4 |  |  |  | 0.31 | 0.04 | 6 |
| E<sup>14</sup>K,S<sup>339</sup>A,S<sup>342</sup>K,D<sup>356</sup>K | 4 |  |  |  | 0.20 | 0.02 | 6 |
| E<sup>14</sup>K,S<sup>339</sup>A,S<sup>342</sup>K,D<sup>352</sup>K,D<sup>356</sup>K | 4 |  |  |  | 0.17 | 0.04 | 6 |

**SI Appendix, Table S2.**
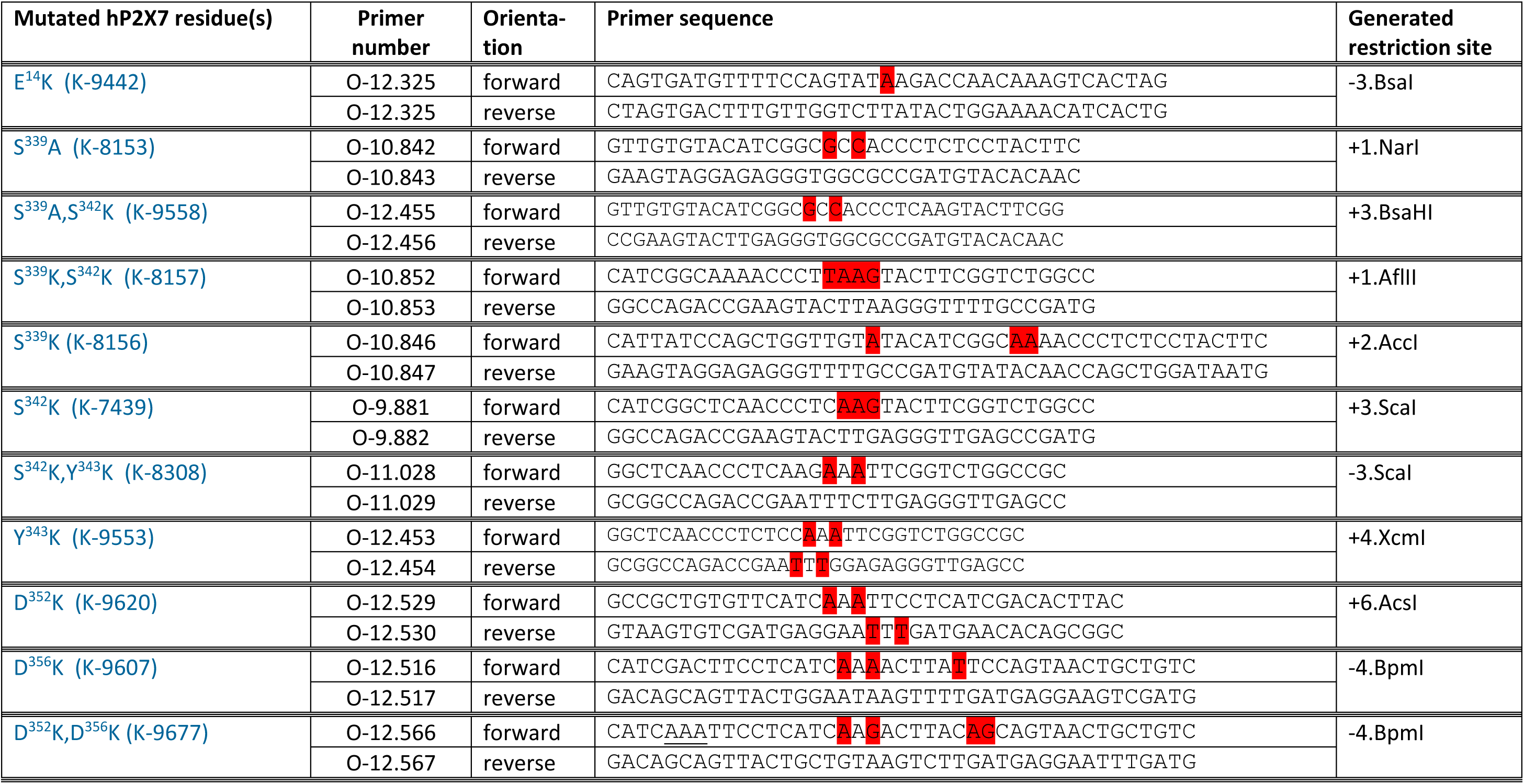
Sequences of mutagenesis primers used in this study. Construct numbers (with ’K-’ prefix) and oligonucleotide numbers (with ’O-’ prefix) refer to internal documentation lists and are used for identification in the laboratory when needed. Nucleotides highlighted in red generate the desired mutation and add or remove the indicated silent restriction site used for initial screening prior to verification by DNA sequencing. The other constructs could be generated from this toolbox of oligonucleotides.

## References

1. S. Schmidt, A. Isaak, A. Junker, Spotlight on P2X7 Receptor PET Imaging: A Bright Target or a Failing Star? International journal of molecular sciences 24 (2023).

2. G. Collo et al., Tissue distribution of the P2X7 receptor. Neuropharmacol 36, 1277–1283 (1997).

3. F. Di Virgilio, A. C. Sarti, S. Falzoni, E. De Marchi, E. Adinolfi, Extracellular ATP and P2 purinergic signalling in the tumour microenvironment. Nat Rev Cancer 18, 601–618 (2018).

4. F. Di Virgilio, G. Schmalzing, F. Markwardt, The elusive P2X7 macropore. Trends Cell Biol 28, 392–404 (2018).

5. A. Surprenant, F. Rassendren, E. Kawashima, R. A. North, G. Buell, The cytolytic P2Z receptor for extracellular ATP identified as a P2X receptor (P2X7). Science 272, 735–738 (1996).

6. P. A. Verhoef, M. Estacion, W. Schilling, G. R. Dubyak, P2X7 receptor-dependent blebbing and the activation of Rho-effector kinases, caspases, and IL-1 beta release. J.Immunol. 170, 5728–5738 (2003).

7. A. Morelli et al., Extracellular ATP causes ROCK I-dependent bleb formation in P2X7-transfected HEK293 cells. Mol.Biol.Cell 14, 2655–2664 (2003).

8. F. Bretschneider, M. Klapperstück, M. Lohn, F. Markwardt, Nonselective cationic currents elicited by extracellular ATP in human and B-lymphocytes. Pflügers Archiv - European Journal of Physiology 429, 691–698 (1995).

9. T. Riedel, I. Lozinsky, G. Schmalzing, F. Markwardt, Kinetics of P2X7 receptor-operated single channels currents. Biophys J 92, 2377–2391 (2007).

10. T. Riedel, G. Schmalzing, F. Markwardt, Influence of extracellular monovalent cations on pore and gating properties of P2X7 receptor-operated single channels currents. Biophys J 93, 846–858 (2007).

11. M. Li, G. E. Toombes, S. D. Silberberg, K. J. Swartz, Physical basis of apparent pore dilation of ATP-activated P2X receptor channels. Nat Neurosci 18, 1577–1583 (2015).

12. M. Harkat et al., On the permeation of large organic cations through the pore of ATP-gated P2X receptors. Proc Natl Acad Sci U S A 114, E3786–E3795 (2017).

13. A. Karasawa, K. Michalski, P. Mikhelzon, T. Kawate, The P2X7 receptor forms a dye-permeable pore independent of its intracellular domain but dependent on membrane lipid composition. eLife 6 (2017).

14. A. E. McCarthy, C. Yoshioka, S. E. Mansoor, Full-length P2X7 structures reveal how palmitoylation prevents channel desensitization. Cell 179, 659–670 e613 (2019).

15. A. Pippel et al., Localization of the gate and selectivity filter of the full-length P2X7 receptor. Proc Natl Acad Sci USA 114, E2156–E2165 (2017).

16. L. Degrève, S. M. Vechi, C. Q. Junior, The hydration structure of the Na^+^ and K^+^ ions and the selectivity of their ionic channels. Biochim Biophys Acta 1274, 149–156 (1996).

17. H. Klein et al., Structural determinants of the closed KCa3.1 channel pore in relation to channel gating: results from a substituted cysteine accessibility analysis. J Gen Physiol 129, 299–315 (2007).

18. X. Bo et al., Pharmacological and biophysical properties of the human P2X5 receptor. Molecular Pharmacology 63, 1407–1416 (2003).

19. I. C. Schiller et al., Dihydropyridines Potentiate ATP-Induced Currents Mediated by the Full-Length Human P2X5 Receptor. Molecules 27 (2022).

20. S. W. Tam, K. Huffer, M. Li, K. J. Swartz, Ion permeation pathway within the internal pore of P2X receptor channels. eLife 12 (2023).

21. A. Nicke et al., P2X1 and P2X3 receptors form stable trimers: a novel structural motif of ligand-gated ion channels. EMBO J 17, 3016–3028 (1998).

22. A. Aschrafi, S. Sadtler, C. Niculescu, J. Rettinger, G. Schmalzing, Trimeric architecture of homomeric P2X2 and heteromeric P2X1+2 receptor subtypes. J Mol Biol 342, 333–343 (2004).

23. D. Becker et al., The P2X7 carboxyl tail is a regulatory module of P2X7 receptor channel activity. J Biol Chem 283, 25725–25734 (2008).

24. M. A. Lomize, I. D. Pogozheva, H. Joo, H. I. Mosberg, A. L. Lomize, OPM database and PPM web server: resources for positioning of proteins in membranes. Nucleic acids research 40, D370–376 (2012).

25. L. Pravda et al., MOLEonline: a web-based tool for analyzing channels, tunnels and pores (2018 update). Nucleic acids research 46, W368–W373 (2018).

26. B. Hille, The permeability of the sodium channel to organic cations in myelinated nerve. J Gen Physiol 58, 599–619 (1971).

27. C. Kubick, G. Schmalzing, F. Markwardt, The effect of anions on the human P2X7 receptor. Biochim Biophys Acta 1808, 2913–2922 (2011).

28. B. N. Cohen, C. Labarca, L. Czyzyk, N. Davidson, H. A. Lester, Tris+/Na+ permeability ratios of nicotinic acetylcholine receptors are reduced by mutations near the intracellular end of the M2 region. J Gen Physiol 99, 545–572 (1992).

29. A. Villarroel, N. Burnashev, B. Sakmann, Dimensions of the narrow portion of a recombinant NMDA receptor channel. Biophys J 68, 866–875 (1995).

30. P. Burow, M. Klapperstück, F. Markwardt, Activation of ATP secretion via volume-regulated anion channels by sphingosine-1-phosphate in RAW macrophages. Pflugers Arch 467, 1215–1226 (2015).

31. F. Markwardt, M. Lohn, T. Böhm, M. Klapperstück, Purinoceptor-operated cationic channels in human B lymphocytes. J.Physiol.(Lond.) 498, 143–151 (1997).

32. T. Kawate, J. C. Michel, W. T. Birdsong, E. Gouaux, Crystal structure of the ATP-gated P2X4 ion channel in the closed state. Nature 460, 592–598 (2009).

33. M. Hattori, E. Gouaux, Molecular mechanism of ATP binding and ion channel activation in P2X receptors. Nature 485, 207–212 (2012).

34. L. Peverini, J. Beudez, K. Dunning, T. Chataigneau, T. Grutter, New Insights Into Permeation of Large Cations Through ATP-Gated P2X Receptors. Frontiers in molecular neuroscience 11, 265 (2018).

35. L. E. Browne et al., P2X receptor channels show threefold symmetry in ionic charge selectivity and unitary conductance. Nat Neurosci 14, 17–18 (2011).

36. C. J. Peters et al., Four basic residues critical for the ion selectivity and pore blocker sensitivity of TMEM16A calcium-activated chloride channels. Proc Natl Acad Sci U S A 112, 3547–3552 (2015).

37. C. Virginio, A. MacKenzie, R. A. North, A. Surprenant, Kinetics of cell lysis, dye uptake and permeability changes in cells expressing the rat P2X7 receptor. J.Physiol (Lond) 519, 335–346 (1999).

38. S. E. Mansoor et al., X-ray structures define human P2X3 receptor gating cycle and antagonist action. Nature 538, 66–71 (2016).

39. L. Cao, H. E. Broomhead, M. T. Young, R. A. North, Polar residues in the second transmembrane domain of the rat P2X2 receptor that affect spontaneous gating, unitary conductance, and rectification. The Journal of neuroscience : the official journal of the Society for Neuroscience 29, 14257–14264 (2009).

40. M. Li, T. H. Chang, S. D. Silberberg, K. J. Swartz, Gating the pore of P2X receptor channels. Nat Neurosci 11, 883–887 (2008).

41. F. Rassendren, G. Buell, A. Newbolt, R. A. North, A. Surprenant, Identification of amino acid residues contributing to the pore of a P2X receptor. EMBO J 16, 3446–3454 (1997).

42. S. Kracun, V. Chaptal, J. Abramson, B. S. Khakh, Gated access to the pore of a P2X receptor: structural implications for closed-open transitions. J Biol Chem 285, 10110–10121 (2010).

43. L. E. Browne, V. Compan, L. Bragg, R. A. North, P2X7 receptor channels allow direct permeation of nanometer-sized dyes. Journal of Neuroscience 33, 3557–3566 (2013).

44. X. Liang, D. S. K. Samways, J. Cox, T. M. Egan, Ca(2+) flux through splice variants of the ATP-gated ionotropic receptor P2X7 is regulated by its cytoplasmic N terminus. J Biol Chem 294, 12521–12533 (2019).

45. A. Nicke et al., A functional P2X7 splice variant with an alternative transmembrane domain 1 escapes gene inactivation in P2X7 knock-out mice. J Biol Chem 284, 25813–25822 (2009).

46. D. S. Samways, B. S. Khakh, S. Dutertre, T. M. Egan, Preferential use of unobstructed lateral portals as the access route to the pore of human ATP-gated ion channels (P2X receptors). Proc Natl Acad Sci U S A 108, 13800–13805 (2011).

47. G. Pierdominici-Sottile, V. Racigh, A. Ormazabal, J. Palma, Charge Discrimination in P2X(4) Receptors Occurs in Two Consecutive Stages. The journal of physical chemistry. B 123, 1017–1025 (2019).

48. V. Racigh, G. Pierdominici-Sottile, J. Palma, Ion Selectivity in P2X Receptors: A Comparison between hP2X3 and zfP2X4. The journal of physical chemistry. B 125, 13385–13393 (2021).

49. M. A. Larkin et al., Clustal W and Clustal X version 2.0. Bioinformatics 23, 2947–2948 (2007).

50. S. Gloor, O. Pongs, G. Schmalzing, A vector for the synthesis of cRNAs encoding Myc epitope-tagged proteins in Xenopus laevis oocytes. Gene 160, 213–217 (1995).

51. M. Klapperstück, C. Büttner, G. Schmalzing, F. Markwardt, Functional evidence of distinct ATP activation sites at the human P2X7 receptor. The Journal of physiology 534, 25–35 (2001).

52. M. P. Weiner et al., Site-directed mutagenesis of double-stranded DNA by the polymerase chain reaction. Gene 151, 119–123 (1994).

53. G. Schmalzing, F. Markwardt, Established Protocols for cRNA Expression and Voltage-Clamp Characterization of the P2X7 Receptor in Xenopus laevis Oocytes. Methods Mol Biol 2510, 157–192 (2022).

54. E. Neher, Ion channels for communication between and within cells. Science 256, 498–502 (1992).

55. W. M. Weber, K. M. Liebold, F. W. Reifarth, W. Clauss, The Ca2+-induced leak current in Xenopus oocytes is indeed mediated through a Cl-channel. The Journal of membrane biology 148, 263–275 (1995).

56. M. Klapperstück, C. Büttner, T. Böhm, G. Schmalzing, F. Markwardt, Characteristics of P2X7 receptors from human B lymphocytes expressed in Xenopus oocytes. Biochim Biophys Acta 1467, 444–456 (2000).

57. M. Klapperstück et al., Antagonism by the suramin analogue NF279 on human P2X1 and P2X7 receptors. Eur J Pharmacol 387, 245–252 (2000).

58. M. Stolz et al., Homodimeric anoctamin-1, but not homodimeric anoctamin-6, is activated by calcium increases mediated by the P2Y1 and P2X7 receptors. Pflugers Arch 467, 2121–2140 (2015).

59. A. L. Lomize, S. C. Todd, I. D. Pogozheva, Spatial arrangement of proteins in planar and curved membranes by PPM 3.0. Protein science : a publication of the Protein Society 31, 209–220 (2022).

60. L. Pauling, The nature of the chemical bond and the structure of molecules and crystals; an introduction to modern structural chemistry, The George Fisher Baker non-resident lectureship in chemistry at Cornell University (Cornell University Press, Ithaca, N.Y.,, ed. 3d, 1960), pp. 644 p.

61. J. Mähler, I. Persson, A study of the hydration of the alkali metal ions in aqueous solution. Inorg Chem 51, 425–438 (2012).

62. C. Virginio, A. MacKenzie, F. A. Rassendren, R. A. North, A. Surprenant, Pore dilation of neuronal P2X receptor channels. Nat Neurosci 2, 315–321 (1999).

63. L. Y. Huang, W. A. Catterall, G. Ehrenstein, Comparison of ionic selectivity of batrachotoxin-activated channels with different tetrodotoxin dissociation constants. J Gen Physiol 73, 839–854 (1979).

64. F. Franciolini, W. Nonner, Anion and cation permeability of a chloride channel in rat hippocampal neurons. J Gen Physiol 90, 453–478 (1987).

65. A. Oyane et al., Formation and growth of clusters in conventional and new kinds of simulated body fluids. J Biomed Mater Res A 64, 339–348 (2003).

## References

1. W. Duckwitz, R. Hausmann, A. Aschrafi, G. Schmalzing, P2X5 subunit assembly requires scaffolding by the second transmembrane domain and a conserved aspartate. J Biol Chem 281, 39561–39572 (2006).

